# Expansion Segments in Bacterial and Archaeal 5S Ribosomal RNAs

**DOI:** 10.1101/2020.08.02.233163

**Authors:** Victor G. Stepanov, George E. Fox

## Abstract

The large ribosomal RNAs of eukaryotes frequently contain expansion sequences that add to the size of the rRNAs but do not affect their overall structural layout and are compatible with major ribosomal function as an mRNA translation machine. The expansion of prokaryotic ribosomal RNAs is much less explored. In order to obtain more insight into the structural variability of these conserved molecules, we herein report the results of a comprehensive search for the expansion sequences in prokaryotic 5S rRNAs. Overall, 89 expanded 5S rRNAs of 15 structural types were identified in 15 archaeal and 36 bacterial genomes. Expansion segments ranging in length from 13 to 109 residues were found to be distributed among 17 insertion sites. The strains harboring the expanded 5S rRNAs belong to the bacterial orders *Clostridiales*, *Halanaerobiales*, *Thermoanaerobacterales*, and *Alteromonadales* as well as the archael order *Halobacterales*. When several copies of 5S rRNA gene are present in a genome, the expanded versions may co-exist with normal 5S rRNA genes. The insertion sequences are typically capable of forming extended helices, which do not seemingly interfere with folding of the conserved core. The expanded 5S rRNAs have largely been overlooked in 5S rRNA databases.

## INTRODUCTION

5S ribosomal RNA (5S rRNA) is a small RNA that is an integral constituent of the central protuberance of the large ribosomal subunit. Although not decisively proven, 5S rRNA is believed to be a mediator of allosteric interactions between the ribosomal functional centers (Dontsova and Dinman 2005, Gongadze 2011, Kiparisov et al. 2005, Kouvela et al. 2007) and an important contributor to efficient and correct ribosome assembly (Huang et al. 2020). On the ribosome, it forms multiple contacts with several structural elements that are linked to the peptidyl transferase center, elongation factor binding region, and mRNA decoding site. Such an intense involvement in intraribosomal interactions imposes severe structural and functional constraints on evolutionary changes, thus making the 5S rRNA one of the most conserved ribosomal components with respect to sequence, secondary structure, and three-dimensional layout [Sun and Caetano-Anolles 2009, Smirnov et al. 2008, Szymanski et al. 1999). The 5S rRNA topology remains essentially the same across all three domains of life. Substantial deviations from the consensus structure are extremely rare and therefore are targets of particular interest.

Expansion segments in 5S rRNAs can certainly be regarded as a very exotic structural abnormality that deserves special attention. The term “expansion segment” is usually applied to taxon-specific nucleotide stretches intervening between the universally conserved elements of large ribosomal RNAs (Weisser and Ban 2019, Parker et al. 2014, Melnikov et al. 2012). Despite being long thought of as a unique distinctive feature of eukaryotic ribosomes, the expansion segments have recently been identified in several bacterial and archaeal 16S and 23S rRNAs (Penev et al. 2020, Armache et al. 2012, Everett et al. 1999, Roller et al. 1992, Greber et al. 2012, Kushwaha and Bhushan 2020, Yang et al. 2017). Meantime, they were never observed in 5S rRNAs apart from two remarkable exceptions. In 1975 Sutton and Woese reported that the 5S rRNA pool from the ribosomes of thermophilic hemicellulolytic bacterium *Thermoanaerobacterium thermosaccharolyticum* (formerly known as *Clostridium thermosaccharolyticum*) consists of two 5S rRNA types, one of the usual length of approximately 120 residues and the other significantly longer containing about 160 nucleotides (Sutton and Woese 1975). After partially assessing RNA sequences using an oligonucleotide fingerprinting assay, they suggested that the oversized 5S rRNA corresponds to an underprocessed precursor with 40 unremoved 3’-terminal residues that is capable to functionally substitute the properly matured 5S rRNA in the ribosomes. They ruled out a possibility that the oversized 5S rRNA represents a novel 5S rRNA species on the grounds that such a drastic change would be very unlikely within the established ribosomal framework made of multiple co-evolved components. However, these conclusions were disproved after the whole genome sequences of several *Th. thermosaccharolyticum* strains became available in the 2010s (Hemme et al. 2010, Jiang et al. 2018, Li et al. 2018). It is now known that this oversized 5S rRNA is encoded separately from the normal 5S rRNA, has the total length of 155 nucleotides, and contains a single 46-nucleotide insertion embedded deep within an otherwise typical 5S rRNA core sequence. In the context of the canonical secondary structure of 5S rRNA, the insertion extends from the universally conserved bulge in helix III. It is predicted to fold as an independent structural module while leaving undisturbed the conventional base pairing scheme of the 5S rRNA core.

In 1981, a second 5S rRNA with an expansion segment was discovered (Luehrsen et al. 1981). It was found that the entire 5S rRNA pool from the ribosomes of a halophilic archaeon *Halococcus morrhuae* consists of molecules far exceeding in size an average archaeal 5S rRNA. Direct enzymatic sequencing of this extra-long 5S rRNA species revealed a 108-nucleotide insertion located immediately adjacent to the so-called “loop E” in the conserved core. The length of the whole molecule was determined to be 231 nucleotides, which is nearly double the usual. Since no other 5S rRNA was found in the *H. morrhuae* cells, this oversized 5S rRNA was assumed to be functional by default. However, the involvement of the expansion segment in ribosomal mechanisms has remained unexplored. Intriguingly, it has been noticed that a closely related archaeon *Halobacterium salinarum* (referred to as *Halobacterium cutirubrum* at the time of the study) has a conventional 5S rRNA without an expansion segment but otherwise highly similar to the *H. morrhuae* 5S rRNA with 89% identity over aligned residues.

Follow-on studies confirmed the presence of expanded 5S rRNAs in all then-known *H. morrhuae* strains and in several uncategorized *Halococcus* isolates (Nicholson 1982). In addition to *H. morrhuae*, the expansion segments were later identified in 5S rRNAs from two other *Halococcus* species, *H. salifodinae* and *H. saccharolyticus* (Stan-Lotter et al. 1999). At the same time, no insertions was found in 5S rRNAs from other haloarchaeal genera, *Halobacterium* and *Haloarcula*, in spite of their close phylogenetic relationships with the genus *Halococcus* (Nicholson 1982, Nicholson and Fox 1983, Daniels et al. 1985).

It is worth noting that the only known 5S rRNAs with expansion segments have been observed in the prokaryotic realm where they would be the least anticipated. This means that both bacterial and archaeal ribosomes are apparently capable of accommodating relatively large additional modules in the immediate vicinity of the 5S rRNA core without losing activity. It does not seem improbable because 5S rRNA is located at the periphery of the ribosome and only partially buried in it. The expansion segments may therefore be rooted in the exposed parts of the 5S rRNA core, and either occupy the nearby cavities or protrude outwards into the surrounding environment (Tirumalai et al. 2020). At the same time, it remains puzzling why so few examples of the expanded 5S rRNAs have been reported so far if the steric constraints on 5S rRNA enlargement are not particularly prohibitive.

In order to assess the abundance of the expanded 5S rRNAs in prokaryotes, we have performed a broad screening of 5S rRNA sequence databases and publicly available genome sequences of bacteria and archaea. A number of novel oversized 5S rRNAs have been identified, some of them harboring not just one but two or even three expansion segments. In the present paper, we describe our search strategy and findings, discuss putative origin and function of the expansion segments in 5S rRNAs, and explain why the expanded 5S rRNAs have been largely overlooked up until now. Throughout the paper, the term “5S rRNA” designates the particular type of ribosomal RNAs rather than refers to the actual size of the RNA itself.

## RESULTS

Description and classification of the expansion segments requires a universal nucleotide numbering system compatible with significant size variations in both bacterial and archaeal 5S rRNAs. For the purpose of the present study, the 5S rRNA bases were numerated according to a previously proposed scheme, which treats the conserved core and each of the insertions as separate nucleotide series (Erdmann and Wolters 1986, Wolters and Erdmann 1988, Fox 1985). Position numbering in the conserved core follows the numbering scheme of the *E. coli* 5S rRNA, which is a long-standing structural archetype of the 5S rRNA family. Positions, which do not belong to the core, are presented in the form of a decimal fraction preceded with a number of the upstream core position as a prefix. Such an approach allows one to accommodate the expansion segments at various insertion points in any 5S rRNA without the necessity of revising nucleotide numbering throughout the entire RNA chain after each new finding (Fig. 1).

**FIGURE 1.**
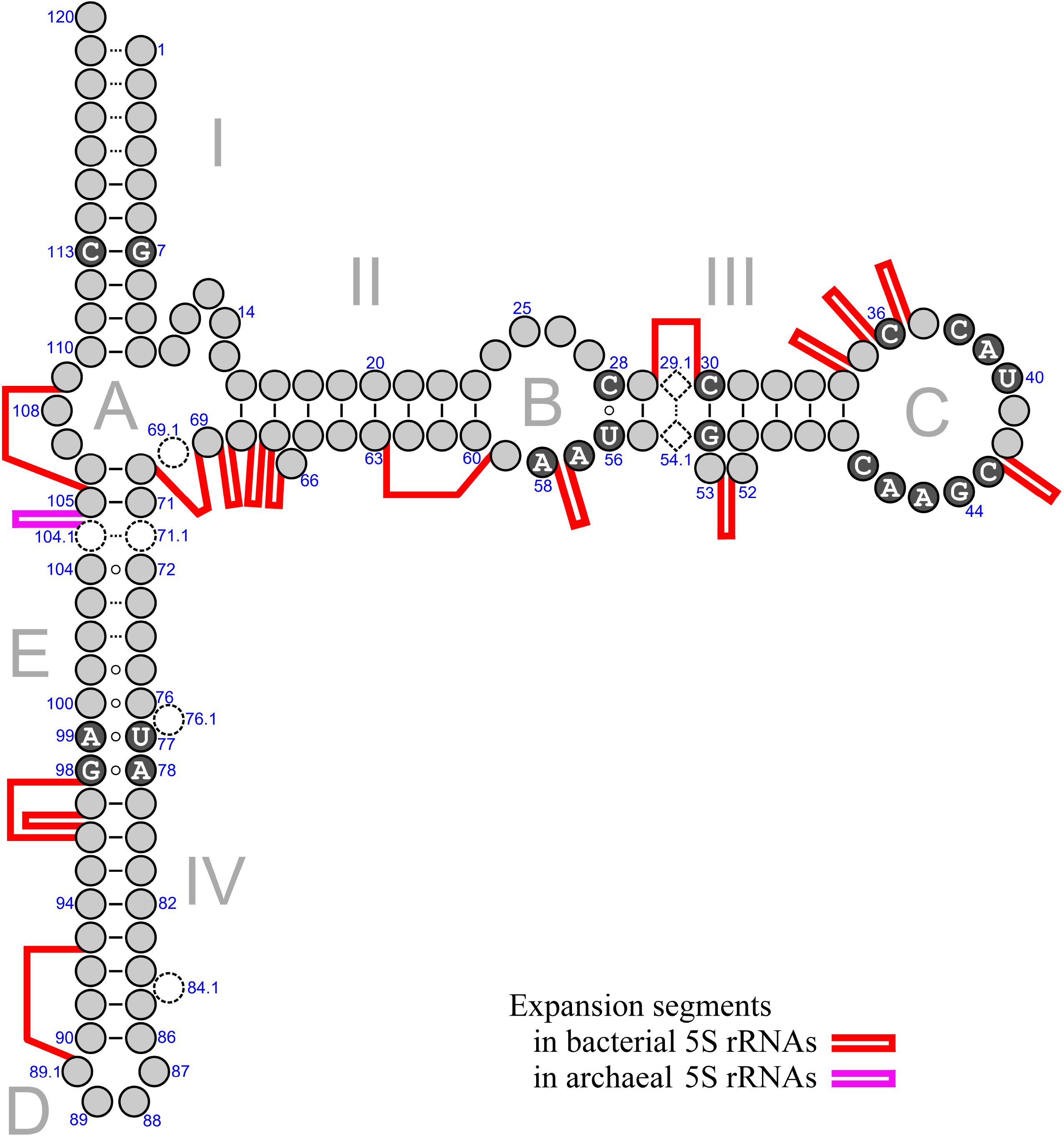
Generalized secondary structure model of prokaryotic 5S rRNA with indicated locations of expansion segments. Universally conserved nucleotides are shown as white characters in black filled circles. Dashed open circles represent nucleotides unique to archaeal 5S rRNAs. Dashed open diamonds represent a unique base pair observed only in 5S rRNAs from the bacterial family *Lachnospiraceae*. Canonical Watson-Crick bonding is depicted with solid lines. Non-Watson-Crick bonding is presented as little open circles. Dotted lines indicate that the bases may be bound by either Watson-Crick or non-Watson-Crick interactions.

Some 5S rRNAs have a shortened Helix I as compared to the *E. coli* 5S rRNA, which makes ambiguous the position numbering in this region. To provide consistent nucleotide nomenclature, we used conserved base pair G7-C113 in Helix I as a landmark. Once the G7-C113 base pair is back tracked from the interior of the molecule, the adjacent residues are numbered relative to this position. Another problematic area is Helix IV, which in some cases is shorter than standard. The solution here was to locate the universal U77-A99 and A78-G98 interactions, and preserve four base Loop D as positions 87, 88, 89, and 89.1 while the stem residues between these two markers were numerated to maximize compatibility with nucleotide conservation pattern observed in Helix IV of regular length. In archaeal 5S rRNAs, base pairs within the “loop E” motif were arranged in order to comply with its layout as seen in the X-ray crystal structure of the *Haloarcula marismortui* 50S ribosomal subunit (Ban et al. 2000, Gabdulkhakov et al. 2013, Szymanski et al. 2002). A single-nucleotide bulge loop was placed between positions 76 and 77 instead of between positions 74 and 75 as in earlier secondary structure models (Wolters and Erdmann 1989, Erdmann and Wolters 1986, Wolters and Erdmann 1988, Erdmann et al. 1987).

The database search for 5S rRNAs with expansion segments initially used the two previously reported examples from the archaeon *H. morrhuae* (Fig. 2A) and the bacterium *Th. thermosaccharolyticum* (Fig. 2B). Both of these 5S rRNAs have one large insertion in the conserved 5S rRNA core. Using these two sequences as queries for BLAST searches of the NCBI *nt* database (ftp://ftp.ncbi.nlm.nih.gov/blast/db/) made it possible to identify similar 5S rRNAs in 15 archaeal strains from the genera *Halococcus*, *Halalkalicoccus*, *Halomarina* and *Halodesulfurarchaeum*, and in 20 bacterial strains from the genera *Thermoanaerobacterium*, *Thermoanaerobacter*, *Caldanaerobacter* and *Caldanaerobius*, respectively. Next, a more rigorous analysis of higher taxons that contain these genera was conducted. A local FASTA database was constructed of 107 complete and draft RefSeq genomes of all strains belonging to the bacterial order *Thermoanaerobacterales*. This database was searched for 5S rRNA-like sequences using RNAMotif software (Macke et al. 2001) with a search pattern that included three 5S rRNA fragments of fixed size corresponding to residues 3-11, 72-80, and 109-117, interspersed with two arbitrary sequences of variable length (50 to 180 nucleotides and 20 to 130 nucleotides, respectively). This search discovered a 172 nucleotide-long 5S rRNA with an expansion segment of novel structural type in *Thermacetogenium phaeum*. A similar search of a database constructed from 257 genomic sequences of all members of the archaeal class *Halobacteria* did not yield any new 5S rRNA sequences with insertions.

**FIGURE 2.**
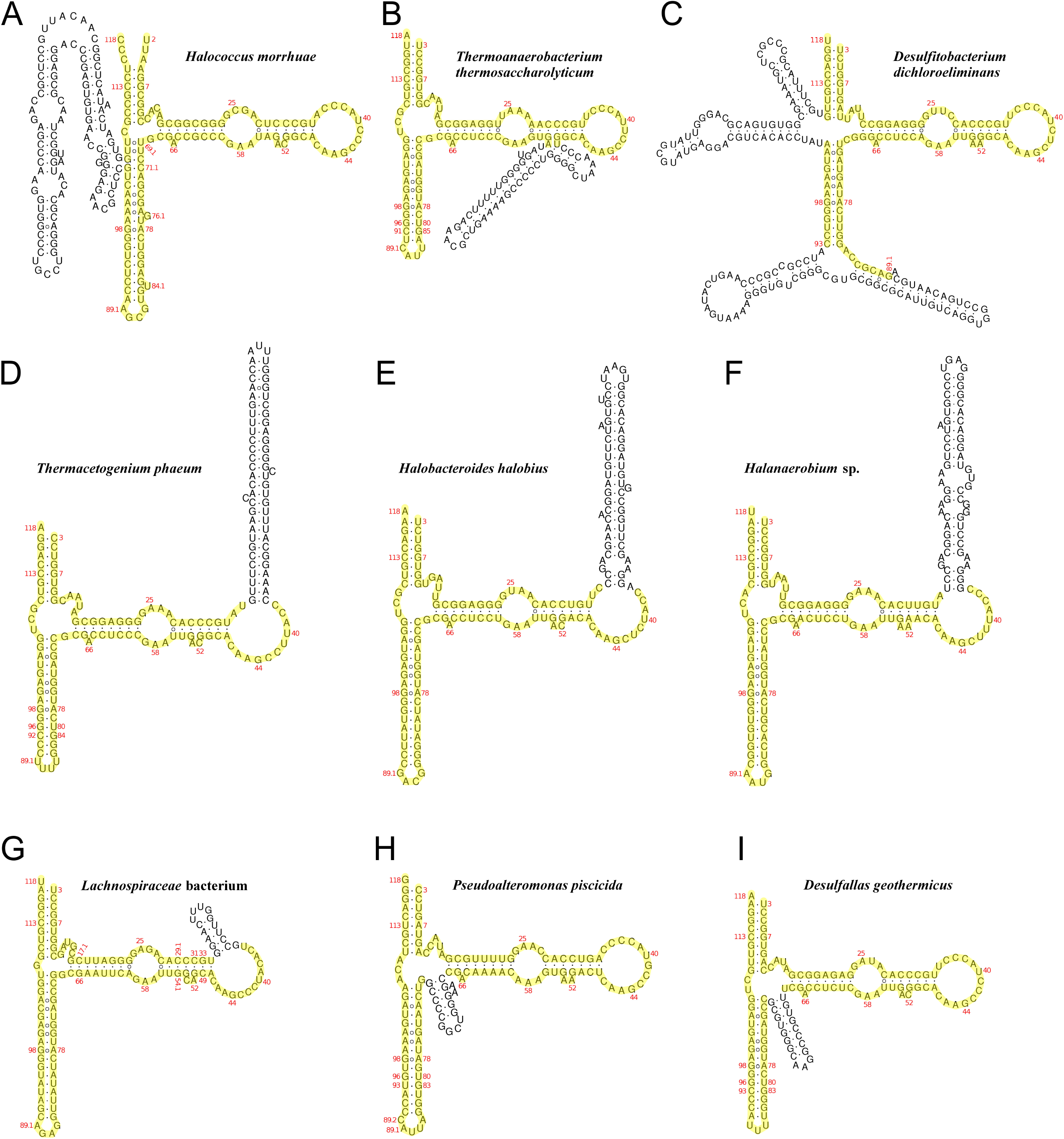
Predicted secondary structures of several representative 5S rRNAs with expansion segments. Nucleotides of the conserved core are highlighted yellow. (*A*) 5S rRNA from *Halococcus morrhuae* DSM 1307 (NZ_AOMC01000054.1, 66311-66541). (*B*) 5S rRNA from *Thermoanaerobacterium thermosaccharolyticum* DSM 571 (NC_014410.1, 84899-85053). (*C*) 5S rRNA from *Desulfitobacterium dichloroeliminans* LMG P-21439 (NC_019903.1, complement (2241573-2241816)). (*D*) 5S rRNA from *Thermacetogenium phaeum* DSM 12270 (NC_018870.1, complement (1414325-1414496)). (*E*) 5S rRNA from *Halobacteroides halobius* DSM 5150 (NC_019978.1, 20186-20360). (*F*) 5S rRNA from *Halanaerobium* sp. DL-01 (NZ_QPJN01000058.1, complement (9-185)). (*G*) 5S rRNA from *Lachnospiraceae* bacterium, strain COE1 (NZ_KE159617.1, 786025-786157). (*H*) 5S rRNA from *Pseudoalteromonas piscicida* DE2-B (NZ_CP021646.1, complement (2103409-2103538)). (*I*) 5S rRNA from *Desulfallas geothermicus* DSM 3669 (NZ_FOYM01000008.1, complement (126967-127097)).

In light of these initial findings, we sought to conduct a more comprehensive search for expanded 5S rRNAs in prokaryotes. Initially, we searched two major collections of 5S rRNA sequences, the 5SrRNAdb database (Szymanski et al. 2016) and the RF00001 family in Rfam database (Kalvari et al. 2018), for sequences with the length of 130 nucleotides and above. In 5SrRNAdb, the search yielded only 6 sequences with the length of 130-132 nucleotides, all of them from prokaryotes. None of these 5S rRNAs had an insertion larger than 2 nucleotides. On the other hand, the RF00001 family of Rfam database was found to contain 1524 RNA sequences within a size range from 130 to a maximum of 346 nucleotides. After removal of sequences with unidentified nucleotides, the number of oversized 5S rRNAs decreased to 1474, with lengths ranging from 130 to 291 nucleotides. Of these, only 34 examples were from prokaryotes (7 from archaea and 27 from bacteria) while the rest mostly represented the 5S rDNA tandems of eukaryotic origin. Each of the selected prokaryotic sequences was evaluated individually by querying the NCBI *nt* database using BLAST, and by aligning it to the closest BLAST hits. This allowed us to identify and exclude from further consideration chimeric sequences and sequences with internal duplication. Of the 16 remaining RNAs, 7 were already known from previous searches, and 9 were newly uncovered 5S rRNAs from the bacterial genera *Desulfallas* and *Desulfofarcimen*, both belonging to the family *Peptococcaceae*, and from an unclassified bacterial genus of the family *Lachnospiraceae*.

In addition to the 5S rRNA database searches, we retrieved all annotated 5S rRNA sequences longer than 129 nucleotides from 1,046 archaeal and 184,876 bacterial RefSeq genomes. In total, 159 oversized 5S rRNA sequences were collected. Of those, 110 sequences had no stretches of unidentified nucleotides, and therefore were retained for deeper analysis. The selected sequences were evaluated for validity as described above, and 52 sequences were discarded because of incorrect boundary assignment or internal sequence duplication. The remaining 58 sequences corresponded to valid 5S rRNAs with insertions. However, the majority of these were already known from the previous searches except for 6 oversized 5S rRNAs from the bacterial genera *Halanaerobium* and *Pseudoalteromonas*.

Finally, we inspected genomes of all organisms carrying oversized 5S rRNAs and their closest relatives for the presence of non-annotated 5S rRNA genes. A local BLAST database was constructed using 478 genomic DNA sequences from the bacterial orders *Clostridiales*, *Halanaerobiales*, *Thermoanaerobacterales*, and *Alteromonadales*, and from the archaeal order *Halobacteriales*. The database was queried with all normal and expanded 5S rRNA sequences available from annotations of the selected genomes. We specifically looked for cases where 5’- and 3’-terminal parts of a query made separate high-scoring hits to a genomic DNA at a distance compatible with the presence of an insertion in the 5S rRNA sequence. This approach allowed us to discover 8 novel expanded 5S rRNAs in *Desulfotomaculum nigrificans*, *D. reducens*, *Desulfitobacterium dichloroeliminans*, and *Halobacteroides halobius* genomes. In the end, the list of the expanded 5S rRNAs contained 89 specimens of 15 structural types found in 15 archaeal and 36 bacterial strains. The number of expansion segments per 5S rRNA molecule varied from 1 to 3, and each of them ranged in length from 13 to 109 nucleotides (Table 1; Supplemental Table S1; Supplemental File S1). In all cases, secondary structure prediction strongly suggested that the expansion segments have extensive internal base pairing, which may allow them to exist as independent modules extending away from an unperturbed 5S rRNA core (Fig. 2A-I; Supplemental Fig. S1-S4).

**TABLE 1.** Occurrence of expanded 5S rRNAs in prokaryotic strains.

| Species | Strain | Expanded 5S rRNAs<br>/Total 5S rRNAs | Expanded 5S rRNA<br>length (nt) | Number of<br>insertions | Insertion length | Insertion position |
| --- | --- | --- | --- | --- | --- | --- |
| *Archaea; Euryarchaeota; Halobacteria; Halobacteriales* |  |  |  |  |  |  |
| *Halococcus morrhuae* | DSM 1307 | 1/1 | 231 | 1 | 108 | 104.1-105 |
| *Halococcus thailandensis* | JCM 13552 | 1/1 | 231 | 1 | 108 | 104.1-105 |
| *Halococcus sediminicola* | CBA1101 | 1/1 | 231 | 1 | 108 | 104.1-105 |
| *Halococcus hamelinensis* | 100A6 | 1/1 | 231 | 1 | 108 | 104.1-105 |
| *Halococcus salifodinae* | DSM 8989 | 1/1 | 232 | 1 | 108 | 104.1-105 |
| *Halococcus saccharolyticus* | DSM 5350 | 1/1 | 232 | 1 | 108 | 104.1-105 |
| *Halococcus agarilyticus* | 197A | 1/1 | 232 | 1 | 108 | 104.1-105 |
| *Halococcus* sp. | IIIV-5B | 1/1 | 231 | 1 | 108 | 104.1-105 |
| *Halalkalicoccus jeotgali* | B3 | 1/1 | 231 | 1 | 108 | 104.1-105 |
| *Halalkalicoccus paucihalophilus* | DSM 24557 | 1/1 | 231 | 1 | 108 | 104.1-105 |
| *Halalkalicoccus* sp. | GSM28 | 1/1 | 231 | 1 | 108 | 104.1-105 |
| *Halomarina oriensis* | SPP-AMP-1 | 1/1 | 228 | 1 | 105 | 104.1-105 |
| *Halomarina oriensis* | JCM 16495 | 1/1 | 228 | 1 | 105 | 104.1-105 |
| *Halodesulfurarchaeum formicicum* | HSR6 | 1/1 | 232 | 1 | 109 | 104.1-105 |
| *Halodesulfurarchaeum formicicum* | HTSR1 | 1/1 | 232 | 1 | 109 | 104.1-105 |
| *Bacteria; Firmicutes; Clostridia; Thermoanaerobacterales* |  |  |  |  |  |  |
| *Thermacetogenium phaeum* | DSM 12270 | 3/3 | 172 | 1 | 61 | 36-37 |
| *Thermoanaerobacterium thermosaccharolyticum* | TG57 | 5/5 | 155 | 1 | 46 | 52-53 |
| *Thermoanaerobacterium thermosaccharolyticum* | DSM 571 | 3/5 | 155 | 1 | 46 | 52-53 |
| *Thermoanaerobacterium thermosaccharolyticum* | M0795 | 4/5 | 155 | 1 | 46 | 52-53 |
| *Thermoanaerobacterium thermosaccharolyticum* | GSU5 | 2/? | 155 | 1 | 46 | 52-53 |
| *Thermoanaerobacterium thermosaccharolyticum* | M5 | 1/? | ? | 1 | ? | 52-53 |
| *Thermoanaerobacterium thermosaccharolyticum* | SP-H2 | 1/? | 155 | 1 | 46 | 52-53 |
| *Thermoanaerobacterium saccharolyticum* | JW/SL-YS485 | 2/5 | 155 | 1 | 46 | 52-53 |
| *Thermoanaerobacterium saccharolyticum* | NTOU1 | 3/5 | 155 | 1 | 46 | 52-53 |
| *Thermoanaerobacterium aotearoense* | SCUT27 | 2/5 | 155 | 1 | 46 | 52-53 |
| *Thermoanaerobacter pseudethanolicus* | ATCC 33223 | 2/4 | 152 | 1 | 43 | 52-53 |
| *Thermoanaerobacter ethanolicus* | CCSD1 | 1/? | 152 | 1 | 43 | 52-53 |
| *Thermoanaerobacter italicus* | Ab9 | 1/4 | 152 | 1 | 43 | 52-53 |
| *Thermoanaerobacter brockii* subsp. *finii* | Ako-1 | 2/4 | 152 | 1 | 43 | 52-53 |
| *Thermoanaerobacter kivui* | DSM 2030 | 3/5 | 152 | 1 | 43 | 52-53 |
| *Thermoanaerobacter* sp. | X513 | 4/4 | 152 | 1 | 43 | 52-53 |
| *Thermoanaerobacter* sp. | X514 | 4/4 | 152 | 1 | 43 | 52-53 |
| *Thermoanaerobacter* sp. | X561 | 3/4 | 152 | 1 | 43 | 52-53 |
| *Thermoanaerobacter* sp. | UBA8867 | 1/? | 152 | 1 | 43 | 52-53 |
| *Caldanaerobacter subterraneus* | DSM 13054 | 1/? | 152 | 1 | 43 | 52-53 |
| *Caldanaerobius polysaccharolyticus* | DSM 13641 | 1/? | 150 | 1 | 41 | 52-53 |
| *Bacteria; Firmicutes; Clostridia; Halanaerobiales* |  |  |  |  |  |  |
| *Halobacteroides halobius* | DSM 5150 | 4/5 | 175 | 1 | 58 | 36-37 |
| *Halanaerobium* sp. | MA284_MarDTE_T2 | 1/? | 177 | 1 | 60 | 36-37 |
| *Halanaerobium* sp. | DL-01 | 1/? | 177 | 1 | 60 | 36-37 |
| *Bacteria; Firmicutes; Clostridia; Clostridiales* |  |  |  |  |  |  |
| *Desulfotomaculum acetoxidans* | DSM 771 | 2/9 | 147; 166 | 1; 2 | 32; 19+32 | 29-30, 42-43 |
| *Desulfallus geothermicus* | DSM 3669 | 1/5 | 131 | 1 | 19 | 69-70 |
| *Desulfallus gibsoniae* | DSM 7213 | 3/9 | 153; 212; 262 | 2; 2; 3 | 19+20; 39+57; 23+39+85 | 42-43, 67-68, 96-98 |
| *Desulfotomaculum nigrificans* | CO-1-SRB | 1/9 | 206 | 1 | 89 | 57-58 |
| *Desulfotomaculum reducens* | MI-1 | 2/10 | 205 | 2 | 13+78 | 59-63, 66-67 |
| *Desulfotomaculum dichloroeliminans* | LMG P-21439 | 1/6 | 244 | 2 | 64+69 | 89.1-93, 105-109 |
| *Lachnospiraceae* bacterium | COE1 | 4/? | 133 | 1 | 15 | 34-35 |
| *Lachnospiraceae* bacterium | A4 | 1/? | 133 | 1 | 15 | 34-35 |
| *Bacteria; Proteobacteria; Gammaproteobacteria; Alteromonadales* |  |  |  |  |  |  |
| *Pseudoalteromonas piscicida* | DE2-A | 1/13 | 130 | 1 | 16 | 68-69 |
| *Pseudoalteromonas piscicida* | DE2-B | 1/13 | 130 | 1 | 16 | 68-69 |
| *Pseudoalteromonas flavipulchra* | NCIMB 2033 | 1/10 | 131 | 1 | 16 | 68-69 |
| *Pseudoalteromonas distincta* | ATCC 700518 | 1/10 | 131 | 1 | 16 | 68-69 |

Question mark, ?, indicates a lack of precise information about 5S rRNA genes due to incompleteness of genome assemblies.

In the course of these searches, several *Peptococcaceae* and *Pseudoalteromonadaceae* genomes were found to contain sequences that exhibit high similarity to 5S rRNA genes except they lack one or both of the 5S rRNA terminal parts and have large insertions in the 5S rRNA-like core (Supplemental File S2). They are usually located downstream of normal 5S rRNA genes at the distal ends of rRNA operons, sometimes in multiple copies, each of which is preceded by a putative transcription terminator. These sequences are likely not being transcribed. However, even if they are transcribed the product RNA likely won’t be processed and folded as a normal 5S rRNA because of the missing parts. Still, they retain enough of the 5S rRNA distinctive features to be identified as former 5S rRNA genes. Species that contain such remnants include *Desulfitobacterium metallireducens*, *Desulfosporosinus orientis*, *D. youngiae*, *Desulfotomaculum nigrificans*, *Desulfallas arcticus*, *Desulfitobacterium dichloroeliminans*, *Pseudoalteromonas piscicida*, *P. distincta*, *P. aliena*, *P. luteoviolacea*, *P. marina*, *P. phenolica*, *P. tetraodonis*, *P. flavipulchra*, and several unclassified strains from the same genera.

Interestingly, some of the insertions in these pseudogenes are nearly identical to the expansion segments of the oversized 5S rRNA genes in the cases when both types are present simultaneously in the same genome. Such a situation was observed in two *Peptococcaceae* strains (*Desulfitobacterium dichloroeliminans* LMG P-21439 and *Desulfotomaculum nigrificans* CO-1-SRB), and three *Pseudoalteromonadaceae* strains (*Pseudoalteromonas piscicida* strains DE2-A and DE2-B, and *P. flavipulchra* ATCC BAA-314). At the same time, several members of these families do not have the expanded 5S rRNA genes but still harbor their pseudogenized counterparts similar to those seen in *D. nigrificans* CO-1-SRB and *P. piscicida* DE2-A /DE2-B genomes, respectively. In two instances, both coming from the genus *Pseudoalteromonas*, the expansion segments embedded in a fragment of 5S rRNA sequence were found on a broad-host-range plasmid at the end of a plasmid-borne ribosomal RNA operon.

The observed resemblance between the obvious pseudogenes and expanded 5S rRNAs raises a question of whether the 5S rRNAs with expansion segments remain functional or have become inactive. There is no doubt about functionality of those expanded 5S rRNAs, which constitute the one and only 5S rRNA type present in a cell, since there is no alternative to them. Such a situation is observed in all species from the archaeal genera *Halococcus*, *Halalkalicoccus*, *Halomarina* and *Halodesulfurarchaeum*, and in the bacterial strains *Thermacetogenium phaeum* DSM 12270, *Thermoanaerobacterium thermosaccharolyticum* TG57, *Thermoanaerobacter* sp. X513 and X514, *Halanaerobium* sp. DL-01 and MA284_MarDTE_T2, and unclassified *Lachnospiraceae* strains COE1 and A4. In all these cases, the expansion segments are expected to not interfere with the 5S rRNA function in the ribosome.

However, when a genome harbors both normal and expanded 5S rRNA genes, it cannot be excluded that the expanded 5S rRNAs are either not expressed or inactive, and only the normal 5S rRNAs are associated with translating ribosomes. In such cases, we sought to assess the functional status of the expanded 5S rRNAs by comparing the immediate genomic context of their coding sequences with that of either normal 5S rRNA genes from the same genome or oversized 5S rRNA genes from closely related organisms harboring only 5S rRNAs of the expanded type. The comparison was performed under the assumption that a loss of function would correlate with accumulation of mutations in the conserved intergenic sites involved in 5S rRNA synthesis and maturation since they would no longer remain under selective pressure. It was found that the majority of the oversized 5S rRNA genes in question can be related to at least one reference gene deemed to produce functional 5S rRNA and embedded within the same genetic context. The strongest indications of functional competence were noted for the expanded 5S rRNAs from the bacterial genera *Thermoanaerobacterium*, *Thermoanaerobacter*, and *Halobacteroides*. However, in several other cases, significant deviations from the conserved layout of prokaryotic 5S rRNA expression units were observed. The most remarkable example is an oversized 5S rRNA gene from *Desulfallas gibsoniae* DSM 7213 (genome accession number NC_021184.1) located on the forward strand at positions 1,812,738 to 1,812,949. It is not a part of an rRNA operon. Instead, it is a stand-alone gene situated between a phosphate transporter pseudogene (DESGI_RS08420) and a sodium-dependent transporter gene with indications of horizontal transfer from a distant species (DESGI_RS08425). While the 5S rRNA coding sequence is preceded by a 107 bp-long region typical for regular 5S rRNA genes in *D. gibsoniae*, it is followed by a transcription terminator, which is not related to any rRNA operon but rather inherited from a protein expression unit similar to those at remote loci DESGI_RS19955 and DESGI_RS20640. Putative promoters are located within the adjacent pseudogene and show little similarity to the promoter groups controlling the expression of rRNA operons. Overall, this 5S rRNA gene looks like a displaced piece of DNA with accidentally acquired transcriptional control elements. As such, it may or may not produce functional 5S rRNA depending on the promoter activity and compatibility of the primary transcript with *D. gibsoniae* 5S rRNA maturation machinery.

Other examples of the oversized 5S rRNA genes with abnormal surroundings include those from *Desulfotomaculum reducens* MI-1 (NC_009253.1, positions 361,111 to 361,315 on the forward strand) embedded in a severely damaged rRNA operon with part of its promoter group lost, *Desulfallas gibsoniae* DSM 7213 (NC_021184.1, positions 1,557,019 to 1,557,280 on the forward strand) with unusual transcription terminator, and *Pseudoalteromonas piscicida* strains DE2-A and DE2-B with multiple substitutions in the promoter group of the corresponding rRNA operon. Also, the oversized 5S rRNA genes from *Pseudoalteromonas flavipulchra* NCIMB 2033, *P. distincta* ATCC 700518, *Caldanaerobacter subterraneus* DSM 13054, and *Caldanaerobius polysaccharolyticus* DSM 13641 were not evaluated due to insufficiency of information for the comparative sequence analysis. For all these problematic cases, the functional state of 5S rRNAs would likely need to be assessed through experimental studies.

It is worth mentioning that the insertional heterogeneity observed in some genomes among 5S rRNA gene copies can also be seen in 16S and 23S rRNA gene families when the expanded rRNA genes co-exist with the normal ones (Supplemental File S3). At the same time, there is no strict correlation between the distribution of large (> 10 bp) insertions in 5S, 16S, and 23S rRNA coding sequences in a single genome with multiple rRNA operons. Of the prokaryotic strains harboring the oversized 5S rRNAs, insertional heterogeneity was detected in *Desulfitobacterium dichloroeliminans* LMG P-21439 5S, 16S and 23S rRNAs, *Desulfofarcimen acetoxidans* DSM 771 5S and 23S rRNAs (but not in 16S rRNAs), *Desulfallas gibsoniae* DSM 7213 5S, 16S and 23S rRNAs, *Thermoanaerobacterium thermosaccharolyticum* DSM 571 5S, 16S and 23S rRNAs, *Th. thermosaccharolyticum* TG57 23S rRNAs (but not in 5S and 16S rRNAs), *Th. thermosaccharolyticum* M0795 5S and 23S rRNAs (but not in 16S rRNAs), *Th. aotearoense* SCUT27 5S and 16S rRNAs (but not in 23S rRNAs), *Thermoanaerobacter brockii* subsp. *finnii* Ako-1 5S, 16S and 23S rRNAs, *Th. pseudethanolicus* ATCC 33223 5S, 16S and 23S rRNAs, and *Th. italicus* Ab9 5S and 23S rRNAs (but not in 16S rRNAs). It seems likely that the mechanisms underlying the insertion of novel modules into the 5S rRNA chain also act on large rRNAs of the mentioned organisms. At the same time, sequences and distribution of the expansion segments in the rRNA gene families point to rather independent acquisition of such modules by 5S, 16S, and 23S rRNA genes.

## DISCUSSION

Our search for the expansion segments in prokaryotic 5S rRNAs revealed 87 novel examples of the expanded 5S rRNA molecules in addition to the previously known two. The Rfam RF00001 and 5SrRNAdb databases and the entire RefSeq collection of annotated bacterial and archaeal genomes have been exhaustively screened for the oversized 5S rRNA molecules. It should be noted that the outcome directly depended on quality of the genome annotations with respect to 5S rRNA gene mapping, and on the content curation practices in the case of 5S rRNA databases. Once identified, the sequences of the expanded 5S rRNAs were used as queries for BLAST similarity search in the NCBI *nt* nucleotide sequence database and whole genome collections. This approach allowed us to discover expanded 5S rRNAs in non- or poorly annotated genomes of species closely related to those from which the queries were derived. Another annotation-independent search method utilized 5S rRNA sequence models, which were constructed of short conserved motifs interspersed with random sequences of variable size, and partly constrained with characteristic secondary structure patterns. It was used to screen only a small fraction of prokaryotic genomes because of high rate of false-positive detections inevitably associated with high redundancy of the search probes. Overall, the most comprehensive search has been performed in the bacterial order *Thermoanaerobacterales* and archaeal class *Halobacteria*, where the initial screening uncovered the majority of novel 5S rRNAs with expansion segments. At the same time, we cannot exclude that non- or poorly annotated genomes outside of these taxonomic groups may still harbor some expanded 5S rRNAs that escaped detection.

One may wonder why the expanded 5S rRNAs were largely overlooked up until now, given that the first two examples have been discovered about four decades ago. One explanation is that substantial number of them have been incorrectly presented as much shorter sequences in genomic annotations and databases in order to make them better fit the consensus-derived standard model of 5S rRNA. For example, all genes encoding the expanded 5S rRNAs in sequenced archaeal genomes have been found misannotated with respect to their boundaries. In all these cases, a fragment of the expansion segment with partial complementarity to the 5’-terminus of 5S rRNA has been misinterpreted as 3’-terminal sequence in order to keep 5S rRNA length within standard range. However, even when correctly mapped in genomic sequences, the expanded 5S rRNAs may remain not deposited in databases and thus evade public attention. The most prominent collection of 5S rRNA sequences, 5SrRNAdb, contains only molecules not longer than 132 residues, possibly because of strict curation practices, which strongly disfavor deposition of anomalous specimens. As a result, the database hosts only those expanded 5S rRNAs, which were erroneously annotated as short enough to match the standard 5S rRNA model. Another specialized database, Rfam RF00001, does not apparently set a rigid size limit for the deposited sequences, and therefore contains several expanded 5S rRNAs with correctly identified boundaries. However, these entries are obscured by numerous artefactual oversized sequences, which originate from sequencing, assembly, and annotation errors.

Interestingly, it has been a chance for some of the expanded 5S rRNAs to be spotted during a study on intragenomic variation of multi-copy 5S rRNA genes in prokaryotes performed in 2012 by Pei *et al* (Pei et al. 2012). Since in a number of cases the expanded 5S rRNAs co-exist with the normals in the same organism, direct comparison of their sequences would immediately expose the presence of expansion segments in the longer molecules. Among the examined genomes, the study mentions those of *Thermoanaerobacter pseudethanolicus*, *Thermoanaerobacter italicus*, *Desulfotomaculum reducens*, and *Desulfotomaculum acetoxidans* (now reclassified as *Desulfofarcimen acetoxidans*), which, as we now know, encode both normal and extended 5S rRNA types. For unclear reasons, the variation between *Th. italicus* 5S rRNA gene copies was not noticed at all, while in the genomes of *Th. pseudethanolicus*, *D. reducens*, and *D. acetoxidans*, the observed differences have been interpreted exclusively in terms of point mutations and single-nucleotide indels. At some point, the authors specifically claim that all the studied 5S rRNA genes have been found to have no “intervening sequences”, i.e. long insertions, as opposed to 16S and 23S rRNA genes. We think that such a conclusion could only be reached if incomplete sequences of the expanded 5S rRNA genes have been used in the study.

In both prokaryotic superkingdoms, the expansion segments are present in only a tiny fraction of 5S rRNAs. At the same time, patterns of their taxonomic distribution within each superkingdom are markedly different. In *Archaea*, the expanded 5S rRNAs are predominantly found within two closely related genera, *Halococcus* and *Halalkalicoccus*, which together form a compact cluster well separated from other members of the family *Halobacteriaceae* (Fig. 3). In every species in this cluster, 5S rRNA has a 108-nucleotide insertion between positions 104.1 and 105 of the archaeal core. The only other clades that harbor the expanded 5S rRNAs are two small genera, *Halodesulfurarchaeum* and *Halomarina*. The recently proposed single-species genus *Halodesulfurarchaeum* is rather remote from the *Halococcus*-*Halalkalicoccus* branch, and grouped with genera *Halobacterium* and *Halarchaeum*, which both are devoid of the expanded 5S rRNAs. Similarly, the genus *Halomarina* belongs to a subfamily whose other members carry only normal 5S rRNA genes. Since all the expansion segments in the oversized 5S rRNAs of the *Halobacteriaceae* family members belong to the same structural type, it is natural to suggest a horizontal transfer of the segment-coding sequence between the *Halococcus*-*Halalkalicoccus* cluster, *Halomarina* and *Halodesulfurarchaeum* genera.

**FIGURE 3.**
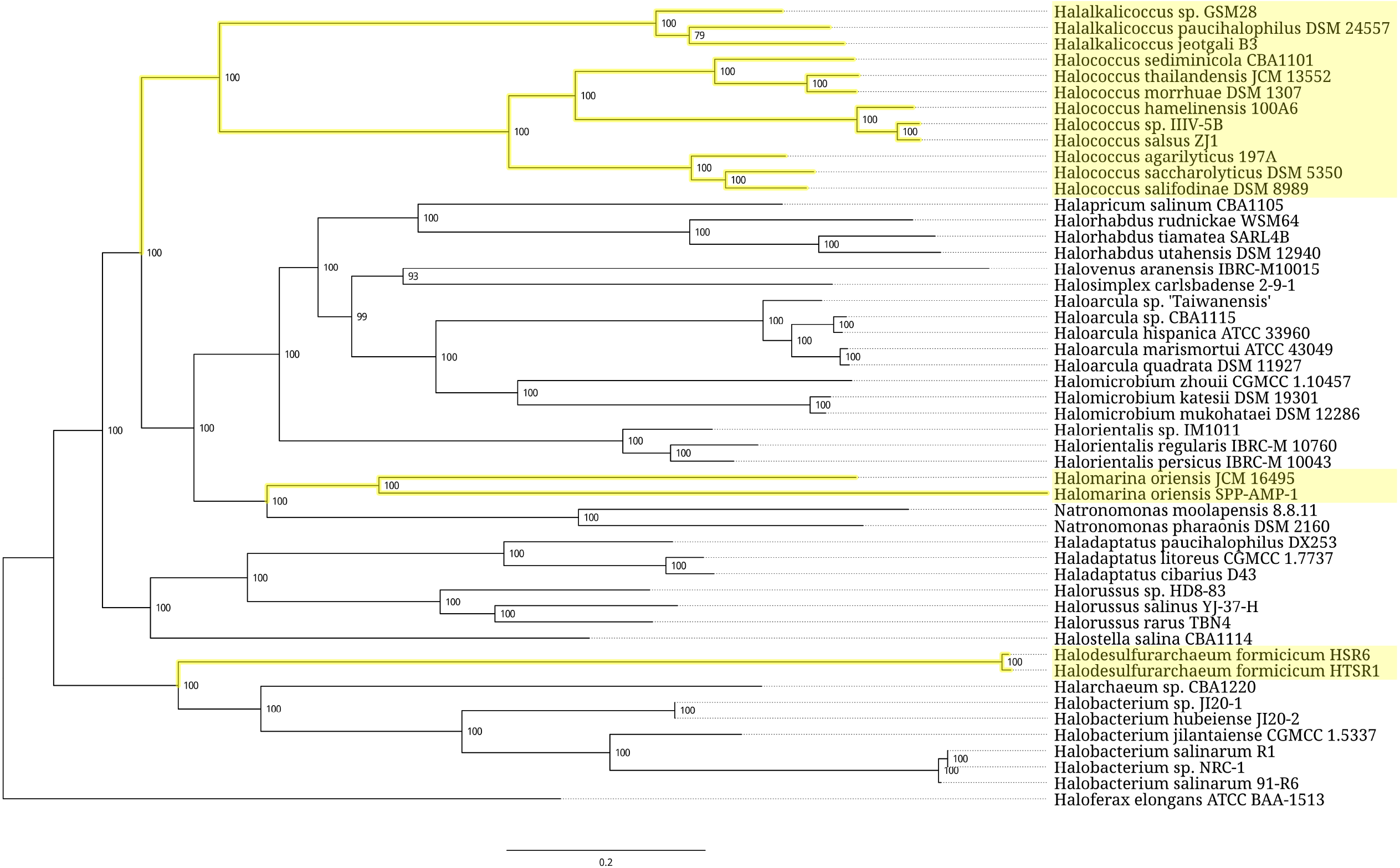
Phylogenetic tree of the archaeal order *Halobacteriales*. Branches associated with the strains carrying the expanded 5S rRNA genes are highlighted yellow. Branch support values represent branch recovery percentage in 100 jackknife resampling replicates. Scale bar represents the number of nucleotide substitutions per position. The tree was generated using the maximum likelihood algorithm on a 106,995 bp-long concatenated codon-wise alignment of 100 conserved single-copy protein-coding genes shared across all of the 50 strains selected for tree construction. The tree was rooted using *Haloferax elongans* ATCC BAA-1513 as an outgroup.

In bacteria, the expansion segments are mostly found in 5S rRNAs of clostridia belonging to the orders *Thermoanaerobacterales*, *Clostridiales*, and *Halanaerobiales*. The only non-clostridial taxon harboring the expanded 5S rRNAs is the gammaproteobacterial genus *Pseudoalteromonas*. The distribution of expanded 5S rRNAs across bacterial taxa is intermittent. This is further complicated by high structural diversity of the expansion modules, which implies their independent origin. It is common for the bacterial species carrying expanded 5S rRNAs to be found together with those with only normal 5S rRNAs within the same genus. Sometime, even strains belonging to the same species have different representation of normal and oversized 5S rRNA types as in the case of *Pseudoalteromonas piscicida* branch. However, the variations in distribution of the expanded 5S rRNA genes seen at subspecies level are not always genuine. They may be caused by strain misassignment as it apparently happened with *Pseudoalteromonas flavipulchra* and *Thermoanaerobacter ethanolicus* lineages.

The most prominent bacterial group associated with the expanded type of 5S rRNA is a cluster of four interconnected thermoanaerobacterial genera, *Thermoanaerobacterium*, *Thermoanaerobacter*, *Caldanaerobacter*, and *Caldanaerobius* (Fig. 4). The group does not look entirely coherent since several branches within it have apparently lost the oversized 5S rRNA genes. All expansion segments within the group belongs to the same structural type, which is represented by a 41-46-nucleotide insertion rooted in the universally conserved bulge at the positions 52 and 53. Alignment of the inserted sequences reveals three distinctive subtypes strictly correlated with phylogenetic branching (Supplemental Fig. 5). One of the subtypes, which originates from the genus *Caldanaerobius*, is apparently ancestral to two others associated with the genus *Thermoanaerobacterium*, and the group *Thermoanaerobacter*-*Caldanaerobacter*, respectively.

**FIGURE 4.**
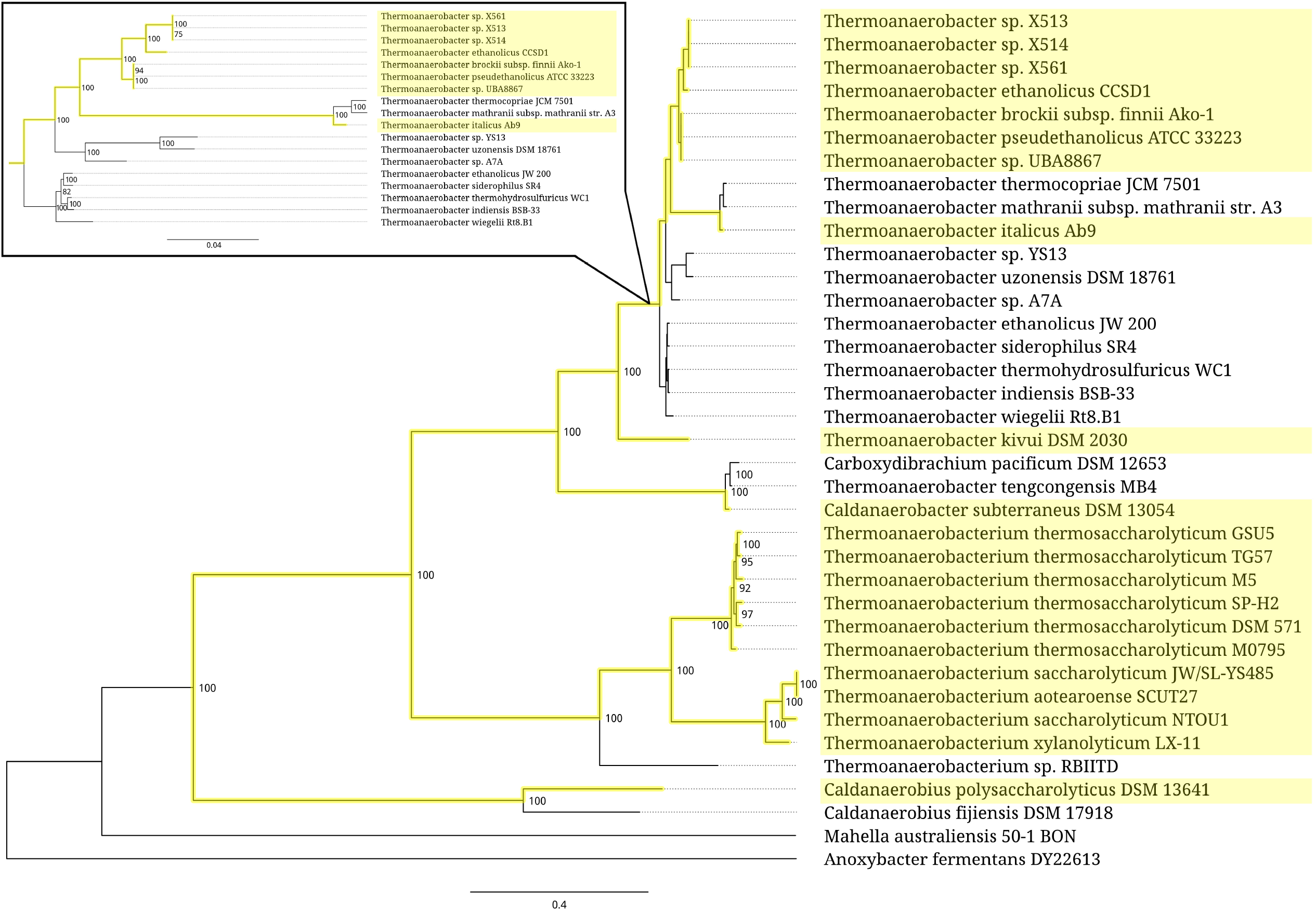
Partial phylogenetic tree of the bacterial order *Thermoanaerobacterales* represented by the related members of families *Thermoanaerobacteraceae*, *Thermoanaerobacterales* Family III and Family IV *incertae sedis*. Branches associated with the strains carrying the expanded 5S rRNA genes are highlighted yellow. The insert shows a scaled-up fragment of the tree. Branch support values represent branch recovery percentage in 100 jackknife resampling replicates. Scale bar represents the number of nucleotide substitutions per position. The tree was generated using the maximum likelihood algorithm on a 80,655 bp-long concatenated codon-wise alignment of 95 conserved single-copy protein-coding genes shared across all of the 37 strains selected for tree construction. The tree was rooted using *Anoxybacter fermentans* DY22613 as an outgroup.

In other cases, the expanded 5S rRNAs are scattered through bacterial taxa as isolated single strains or small clusters consisting of several closely related strains (Fig. 5-7). At the above-species levels, the expansion sequences may propagate via horizontal transfer apparently mediated by mobile genetic elements. One such a long-range horizontal link connects the *Halobacteroides halobius* DSM 5150 strain from the family *Halobacteroidaceae* with a group of two unclassified *Halanaerobium* strains, DL-01 and MA284_MarDTE_T2, from the family *Halanaerobiaceae* (Fig. 6). Another horizontal transfer of the expansion sequence seemingly happened between a group of two *Pseudoalteromonas piscicida* strains, DE2-A and DE2-B, and a group consisting of *P. distincta* ATCC 700518 and *P. flavipulchra* ATCC BAA-314 strains (Fig. 7). It should be noted, however, that multiple occurrences of 5S rRNA remnants with the same insertion as in expanded 5S rRNAs hinder precise identification of a donor of the transferred sequence among members of the family *Pseudoalteromonadaceae*. In addition to the horizontal route, acquisition of the expansion segments might also proceed through vertical inheritance of the pseudogenes followed by their homologous recombination with normal 5S rRNA genes. Similar considerations can be applied to the distribution pattern of the expanded 5S rRNAs in the family *Peptococcaceae*, whose members also occasionally harbor the 5S rRNA gene remnants with insertions.

**FIGURE 5.**
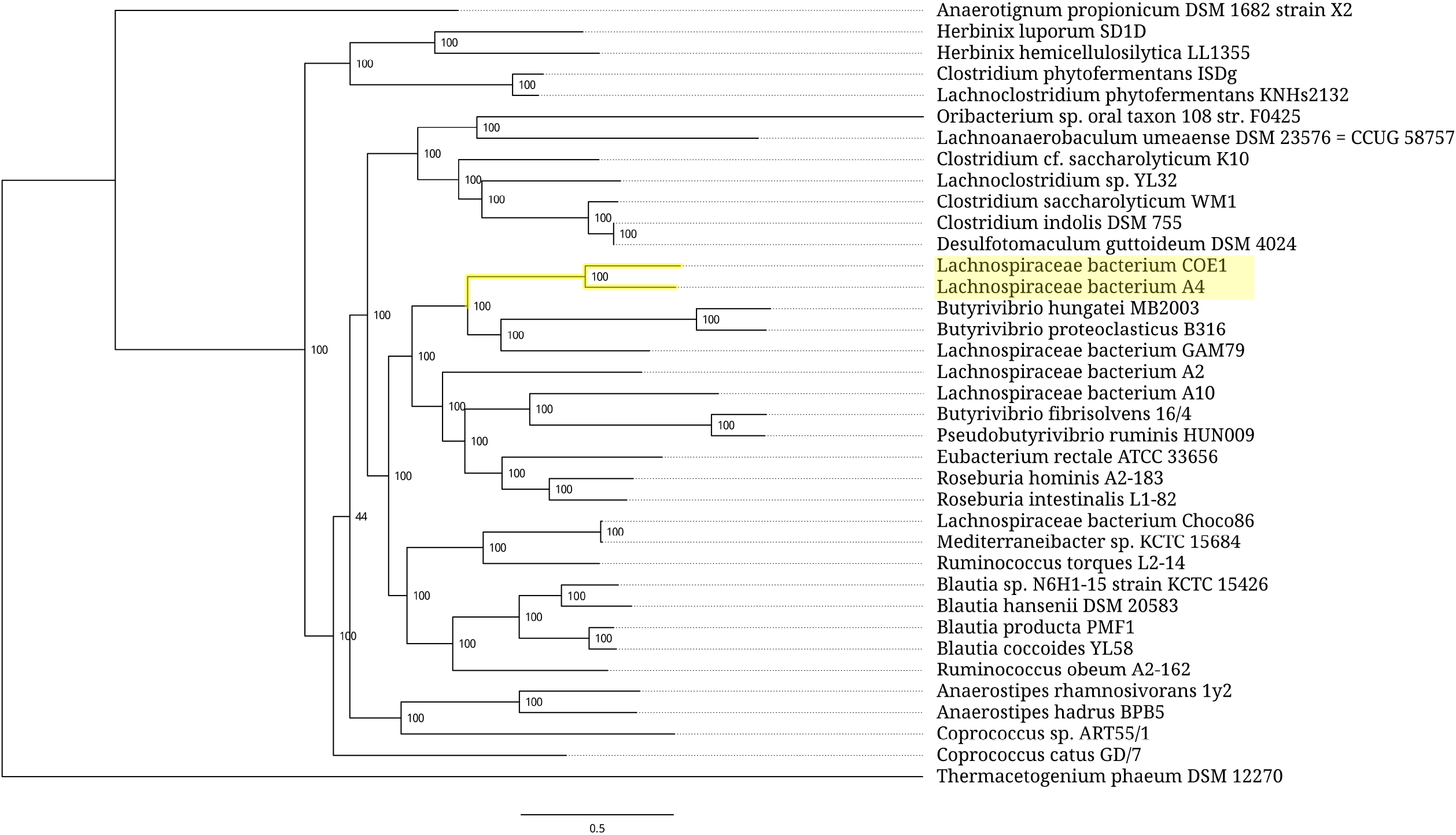
Phylogenetic tree of the bacterial family *Lachnospiraceae*. Branches associated with the strains carrying the expanded 5S rRNA genes are highlighted yellow. Branch support values represent branch recovery percentage in 100 jackknife resampling replicates. Scale bar represents the number of nucleotide substitutions per position. The tree was generated using the maximum likelihood algorithm on a 98,322 bp-long concatenated codon-wise alignment of 100 conserved single-copy protein-coding genes shared across all of the 37 strains selected for tree construction. The tree was rooted using *Thermacetogenium phaeum* DSM 12270 as an outgroup.

**FIGURE 6.**
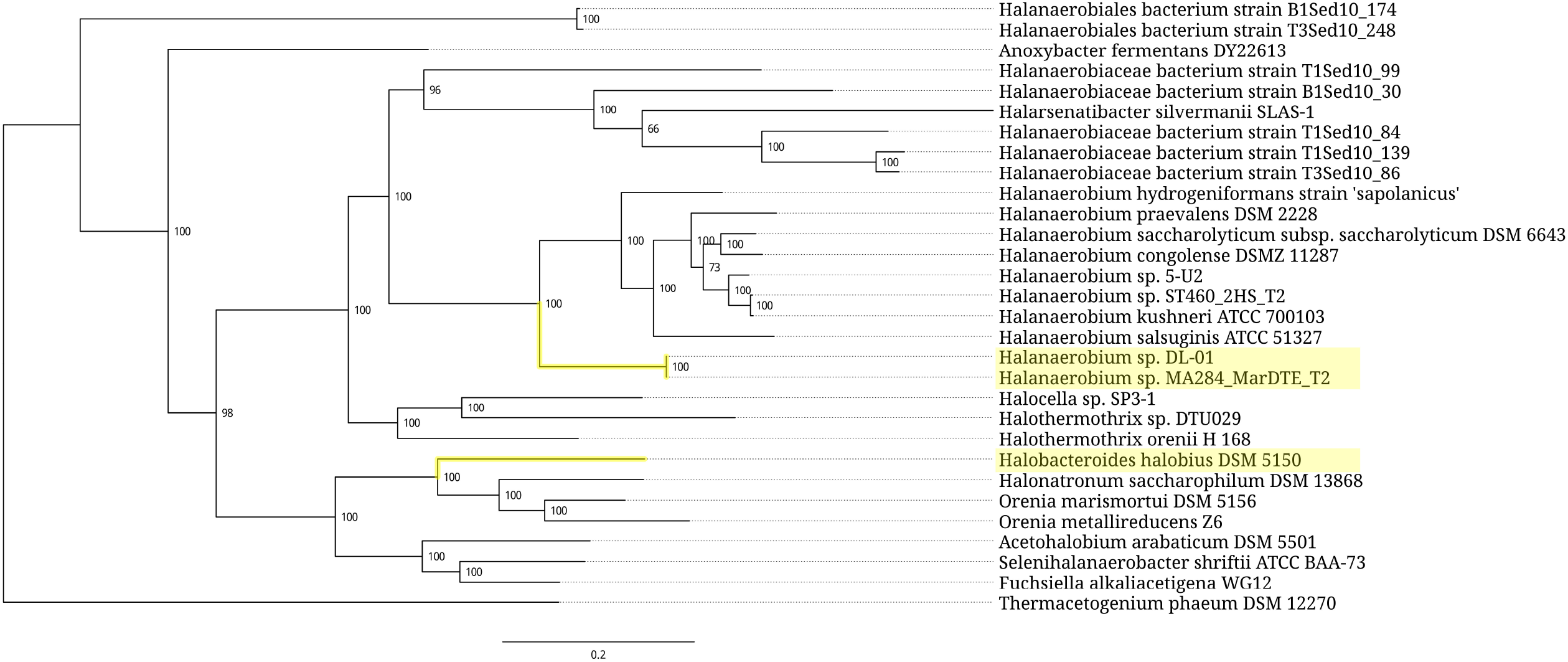
Phylogenetic tree of the bacterial order *Halanaerobiales*. Branches associated with the strains carrying the expanded 5S rRNA genes are highlighted yellow. Branch support values represent branch recovery percentage in 100 jackknife resampling replicates. Scale bar represents the number of nucleotide substitutions per position. The tree was generated using the maximum likelihood algorithm on a 29,271 bp-long concatenated codon-wise alignment of 27 conserved single-copy protein-coding genes shared across all of the 30 strains selected for tree construction. The tree was rooted using *Thermacetogenium phaeum* DSM 12270 as an outgroup.

**FIGURE 7.**
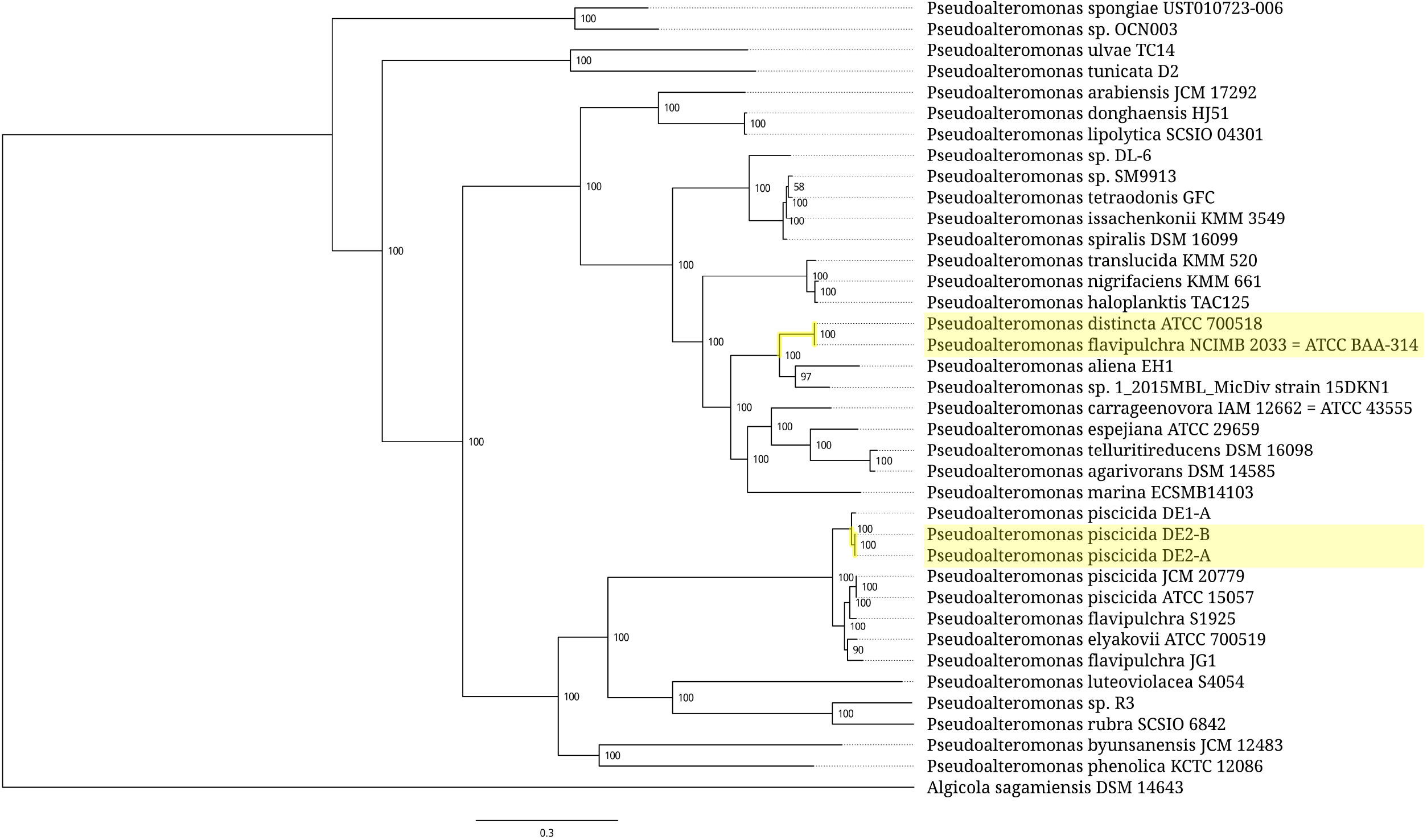
Phylogenetic tree of the bacterial genus *Pseudoalteromonas*. Branches associated with the strains carrying the expanded 5S rRNA genes are highlighted yellow. Branch support values represent branch recovery percentage in 100 jackknife resampling replicates. Scale bar represents the number of nucleotide substitutions per position. The tree was generated using the maximum likelihood algorithm on a 126,927 bp-long concatenated codon-wise alignment of 100 conserved single-copy protein-coding genes shared across all of the 38 strains selected for tree construction. The tree was rooted using *Algicola sagamiensis* DSM 14643 as an outgroup.

Unlike their counterparts in eukaryotic 18S and 25S/28S rRNAs, the expansion segments in prokaryotic 5S rRNAs do not follow any recognizable trend that correlates with phylogenetic branching order across the bacterial and archaeal superkingdoms. Instead, they are mostly divided in taxonomically remote and structurally unrelated groups, each of which represents a rather uniform set of sequences. The non-contiguous taxonomic distribution alongside with rare occurrence point to a non-universal character of their adaptive value, if any exists. Therefore, even if an expansion of 5S rRNA contributes to the fitness of host organism, this contribution is likely restricted to the conditions uncommon for the majority of prokaryotes. Otherwise it would be difficult to explain why the expanded 5S rRNAs have not spread through the entire prokaryotic realm but instead remain confined within boundaries of several particular taxa of subfamily ranks.

Analysis of environmental preferences of the prokaryotic hosts shows a correlation between the presence of expanded 5S rRNA genes and microaerobic/anaerobic lifestyle accompanied with strong halophilic or thermophilic traits. Thus, the insertions in 5S rRNAs might, for example, be associated with peculiar metabolic pathways involved in niche-specific adaptation to hypoxic saline or thermal habitats. At the same time, we cannot completely rule out a possibility that some of the expansion segments do not confer any adaptive benefit for the hosts, and instead represent accidentally acquired features undergoing neutral evolution. While this may not seem likely in the case of relatively large and genetically cohesive taxonomic clusters, such as *Halococcus-Halalkalicoccus* and *Thermoanaerobacterium*-*Thermoanaerobacter*-*Caldanaerobacter*-*Caldanaerobius* groups, it is certainly possible for the small and incoherent ones, like the set of strains from the family *Peptococcaceae*.

The observation that bacterial genomes can host both normal and expanded 5S rRNA genes leads to interesting consequences in the form of heterogeneity of the translation apparatus. The two types of 5S rRNA must confer different operational specializations on the ribosome in order to simultaneously persist within the same organism over an evolutionary significant period of time. Thus, the ribosomal population would consist of two subpopulations optimized for different tasks. In the cases when 16S and 23S rRNAs are also represented by several different structural types, the number of possible rRNA combinations would increase accordingly, giving rise to even more diversified ribosomal pool. Such a structural heterogeneity of the translation apparatus is anticipated to increase its functional flexibility. It would allow cells to finely adjust the translation efficiency and accuracy to current metabolic requirements through differential expression of rRNA genes of various types. Up until now, the concept of ribosomal heterogeneity has mostly been associated with variations in protein composition of eukaryotic ribosomes (Genuth and Barna 2018a, Genuth and Barna 2018b, Ferretti and Karbstein 2019, Byrgazov et al. 2013, Sauert et al. 2015). The present study implies that ribosomal heterogeneity can also be achieved in certain prokaryotes via combinatorial assembly of the ribosomes from a plethora of normal and expanded rRNA types.

The distribution of expansion segments across the elements of prokaryotic 5S rRNA secondary structure model is expectedly uneven (Fig. 1). The insertion sites are concentrated in the loop B - helix III - loop C fragment, in the vicinity of the triple helical junction, and in the 3’-terminal half of the helix IV. The inserted modules are anticipated to not interfere with folding of the 5S rRNA core because of their extensive internal base pairing and lack of noticeable complementarity to the core sequence. Also, the insertions rarely displace bases of the core or disrupt tertiary interactions, which contribute to maintenance of the core structure. At the same time, not all of the insertions seem compatible with the immediate surroundings of 5S rRNA on the ribosome. In archaeal 5S rRNAs, the sole identified insertion site (pos. 104.1-105) allows unhindered sequence expansion away from other ribosomal constituents (Fig. 8A). In bacteria, the insertion site at positions 29-30 overlaps with the binding site of ribosomal protein uL18 while positions 42-43 and 57-58 are blocked by protein uL5. In addition, substantial steric constraints disfavor a large sequence expansion at positions 68-70 and 89.1-93 whereas new modules rooted at positions 66-68 and 96-98 would encounter significantly milder spatial restrictions. The least obstructed insertion sites in bacterial 5S rRNAs are located at positions 34-37 in loop C, and positions 52-53 corresponding to the universally conserved bulge in helix III (Fig. 8B).

**FIGURE 8.**
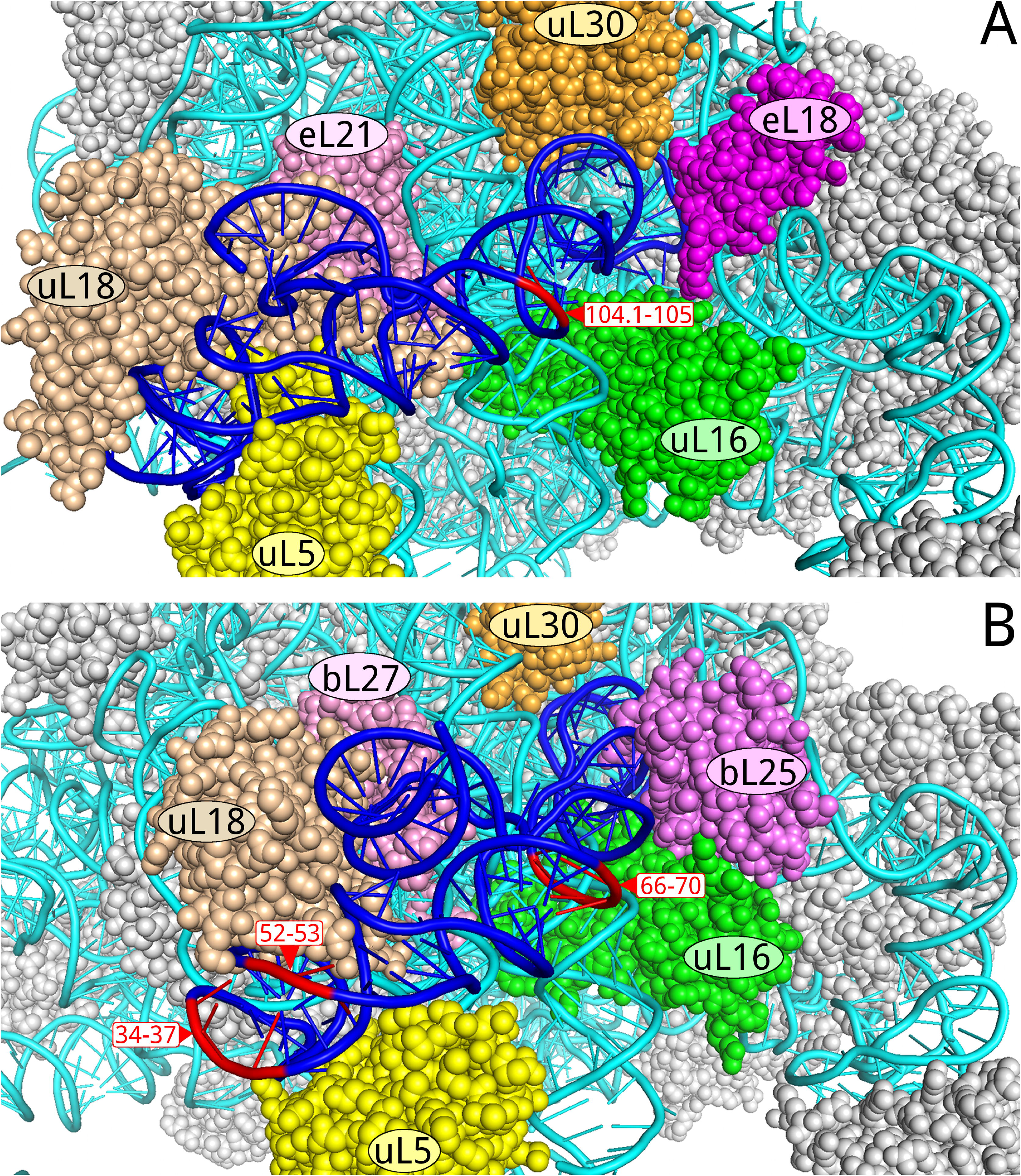
Apical part of large ribosomal subunit viewed from solvent side. Ribosomal proteins are presented in space-filling mode, rRNAs are shown as pipes. 5S rRNA is colored blue except for the rooting sites of the most exposed expansion segments, which are shown in red. 23S rRNA is colored cyan. (*A*) Archaeal 50S subunit (*Haloarcula marismortui*, PDB 4V9F). (*B*) Bacterial 50S subunit (*Escherichia coli*, PDB 4YBB).

Taking these considerations into account, one might conclude that the expanded 5S rRNAs from *Desulfofarcimen acetoxidans*, *Desulfallas gibsoniae*, and *Desulfotomaculum nigrificans* cannot properly bind to the large ribosomal subunit because their interactions with associated ribosomal proteins uL5 and uL18 are compromised. Unless the expansion segments can functionally substitute for these proteins, these 5S rRNAs are unlikely to correctly assemble with other ribosomal components to produce functional ribosome. Likewise, a 69-nucleotide insertion in the helix IV of the expanded 5S rRNA from *Desulfitobacterium dichloroeliminans* may prevent binding of the whole molecule to the ribosome due to steric constraints at the helix IV docking cavity. In all these cases, the expansion of the 5S rRNA chain possibly represents the initial stage of pseudogenization or a shift to a novel extraribosomal function.

The most conspicuous types of expansion segments are predictably associated with the three most exposed insertion sites. In halobacterial 5S rRNAs, the expansion segment is located between the “loop E” and triple helical junction, and expected to extend outwards of the ribosome. It is shaped as an asymmetric “V” with the shorter arm folded as a hairpin, and longer arm as an unbranched protrusion consisting of several helical stems interspersed with internal loops (Fig. 2A, Supplemental Fig. S6). The distal stem-loop element is the most conserved part of the insertion, and therefore a prime candidate for a putative functional domain.

Expansion of thermoanaerobacterial 5S rRNAs predominantly occurs at the conserved bulge in helix III in a form of two hairpins of unequal size (Fig. 2B, Supplemental Fig. S7). The entire fragment is situated at the very top of the central protuberance. The longer hairpin is homoiteron-rich, which is a feature often seen in the expansion segments of eukaryotic rRNAs (Parker et al. 2015). Interestingly, similar motifs are present in several prokaryotic genomes other than thermoanaerobacterial. They reside mostly in intergenic regions, including the leaders and spacers in rRNA operons. Their possible functions inferred from their locations include transcription control and RNA processing. It is important to note that in the thermoanaerobacterial genomes, such motifs are found only in the expanded 5S rRNA genes.

The insertion sites at positions 34-37 in the loop C harbor expansion segments in 5S rRNAs from several distant clostridial strains belonging to the families *Thermoanaerobacteraceae*, *Halobacteroidaceae*, *Halanaerobiaceae*, and *Lachnospiraceae*. Apart from the latter case, the inserted modules represent long unbranched helices with bulges and internal loops. In this respect, they bear a superficial resemblance to the long arm of the expansion segment from halobacterial 5S rRNAs. Despite being in close proximity to the binding sites of proteins uL5 and uL18, the inserted helices have enough space for accommodation without any interference with the nearby ribosomal constituents.

Seemingly unhindered accommodation of the long helical fragments inserted at positions 34-37, 52-53, and 104.1-105 highlights a potential value of these sites for biotechnological applications. A regular prokaryotic 5S rRNA could be modified to harbor artificial RNA modules at these locations in order to confer novel properties and functions on the ribosome. The modules may carry aptamers, ribozymes, riboswitches, antisense sequences, purification tags and other RNAs of interest. Genetically modified 5S rRNA might be supplemented with altered 16S and 23S rRNAs if multiple artificial modules have to be introduced into a ribosomal particle. Mobile expression constructs with modified 5S rRNA genes might also be used to create and control multiple ribosomal subpopulations in a cell with its own set of natural 5S rRNA genes.

While being commonly regarded as a molecule with highly conserved structure, 5S rRNA can apparently accommodate relatively large expansion segments comparable in size with its core. This finding gives us a better understanding of structural constraints imposed on 5S rRNA by the surrounding ribosomal framework. It also provides us with new insight into admissible variations in the prokaryotic translation apparatus. The acquired knowledge could be useful in studies on function and evolution of 5S rRNA, and in ribosome redesign for biotechnological applications.

## MATERIALS AND METHODS

### Databases

The nucleotide *nt* database was downloaded from the National Center for Biotechnology Information (NCBI) FTP site ftp://ftp.ncbi.nlm.nih.gov/blast/db/ in April 2020, and used for local BLAST searches. Complete and draft RefSeq assemblies of all available archaeal and bacterial genomes were downloaded from the NCBI FTP site ftp://ftp.ncbi.nlm.nih.gov/genomes/all/GCF/ on April 25, 2018, using the corresponding assembly_summary.txt files as a source of ftp paths to the assembly directories. A homemade Perl script was used to retrieve 5S rRNA sequences with minimum size limit of 130 nucleotides from the [accession number]_[assembly name]_rna_from_genomic.fna.gz files. When necessary, local BLAST or FASTA databases were created from genomic DNA sequences of the selected assemblies. 5SrRNAdb (Szymanski et al. 2016) and Rfam RF00001 (Kalvari et al. 2018) collections of 5S rRNA sequences were last accessed on May 4, 2020, at http://combio.pl/rrna/ and http://rfam.xfam.org/family/RF00001/, respectively.

### Sequence analysis

Searches of the genomic nucleotide databases for 5S rRNA-coding sequences were performed using the Basic Local Alignment Search Tool (BLAST) 2.7.1+ (Camacho et al. 2009). Handling and analysis of short DNA/RNA sequences was managed using Seaview 4.6.1 software package (Gouy et al. 2010). Annotated genomes were explored using the Artemis genome viewer 17.0.1 (Rutherford et al. 2000, Carver et al. 2008).

A 5S rRNA pattern matching search of the *Thermoanaerobacterales* and *Halobacteria* genomes was performed using RNAMotif 3.1.1 (Macke et al. 2001). Descriptors for the 5S rRNA-like secondary structure pattern were constructed to allow large insertions anywhere in the structure except helix I and the upstream half of “loop E”. In the case of *Thermoanaerobacterales*, the search pattern was YYYGGYGRYN_50-180_GAUGRUASUN_20-150_YYGCCRRR, with the 5’-terminal fragment YYYGGYGR required to be complementary to the 3’-terminal fragment YYGCCRRR. In the case of *Halobacteria*, the search pattern was MGGCGGCCN_50-180_BNNMSUACUN_20-150_TCGCCGCC, where the 5’-terminal fragment GGCGGC was base-paired with the 3’-terminal fragment GCCGCC.

The identified 5S rRNA expansion segments were folded as self-sufficient structural modules using an energy-based algorithm from RNAstructure 6.0 (Xu and Mathews 2016). RNA secondary structure prediction driven by both energy constraints and sequence covariation data was performed using R-scape 0.3.3 (Rivas et al. 2017) and the RNAalifold program from the ViennaRNA 2.2.5 software package (Lorenz et al. 2016, Hofacker and Lorenz 2014). 5S rRNA secondary structures were visualized using RNAviz 2.0.3 (De Rijk et al. 2003).

### Phylogenetic tree construction

Phylogenetic trees were generated using tools provided at the Pathosystems Resource Integration Center (PATRIC) website, https://patricbrc.org (Wattam et al. 2014, Wattam et al. 2017). The trees were inferred from concatenated multiple alignments of the conserved single-copy genes of homologous proteins using the Codon Tree method. Typically, up to 100 of shared proteins were selected specifically for each group of prokaryotic genomes under analysis. The combined protein/coding DNA sequence alignments were processed using the program RAxML for maximum likelihood tree building (Stamatakis 2014). Robustness of the built trees was assessed using the gene-wise jackknifing procedure with 100 resampling replicates per tree. The trees were visualized using FigTree software (http://tree.bio.ed.ac.uk/software/figtree/).

## SUPPLEMENTAL MATERIAL

Supplemental material is available for this article.

## ACKNOWLEDGMENTS

This work was supported in part by NASA Exobiology Grant NNX14AK36G and NASA Contract 80NSSC18K1139 under the Center for Origin of Life, Georgia Institute of Technology to GEF.

