## Supplemental Fig. S1 for "Expansion Segments in Bacterial and Archaeal 5S Ribosomal RNAs"

**Desulfallas gibsoniae 5S rRNA**
**NC_021184.1[1557019-1557280]fwd**

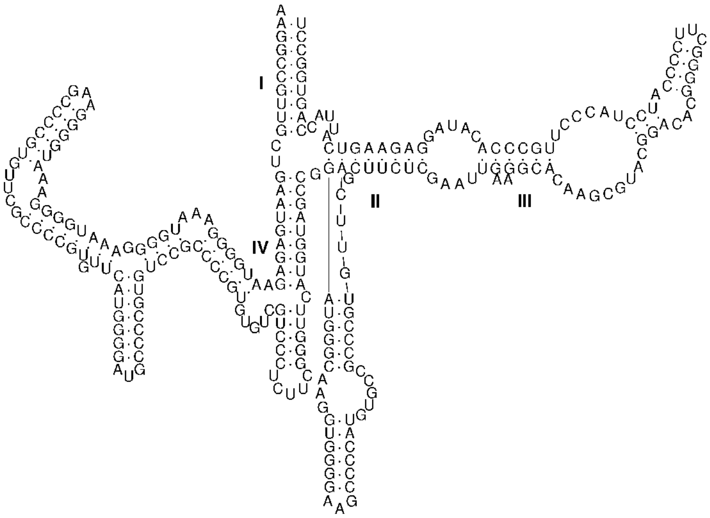


**Desulfallas gibsoniae 5S rRNA**
**NC_021184.1[1760679-1760831]fwd**

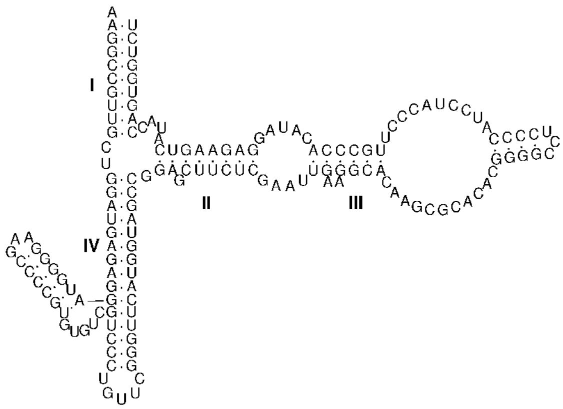


**Desulfallas gibsoniae 5S rRNA**
**NC_021184.1[1812738-1812949]fwd**

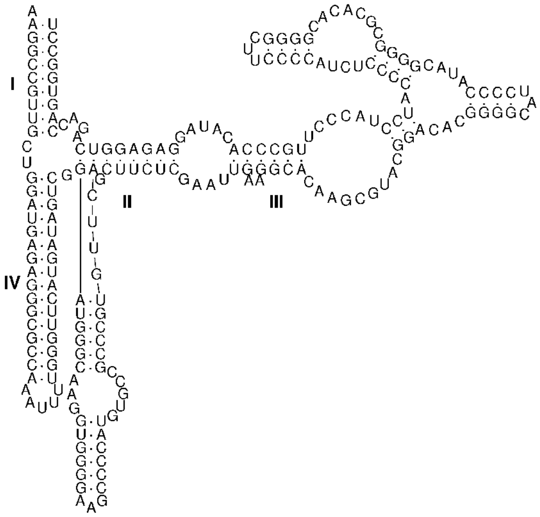


**Desulfallas gibsoniae 5S rRNA**
**NC_021184.1[14594-14708]fwd**

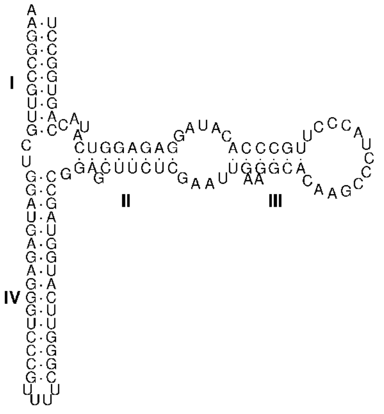


**Supplementary Figure 1**. Variant 5S rRNAs with or without expansion segments identified in *Desulfallus gibsoniae* DSM 7213 genome.
