## Supplemental Fig. S2 for "Expansion Segments in Bacterial and Archaeal 5S Ribosomal RNAs"

**Desulfofarcimen acetoxidans 5S rRNA**
**NC_013216.1[2045104-2045250]fwd**

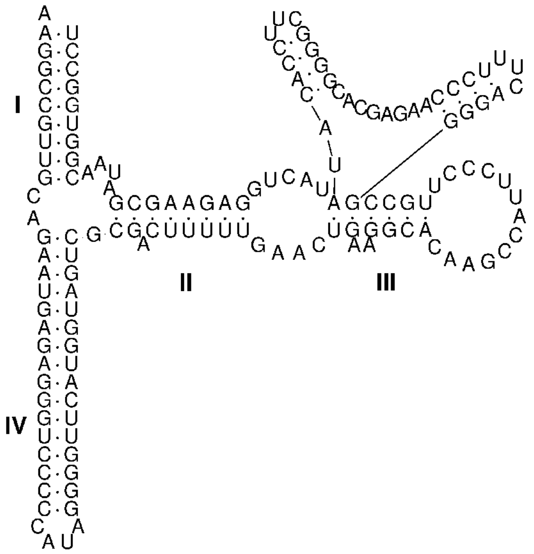


**Desulfofarcimen acetoxidans 5S rRNA**
**NC_013216.1[2945859-2946024]rev**

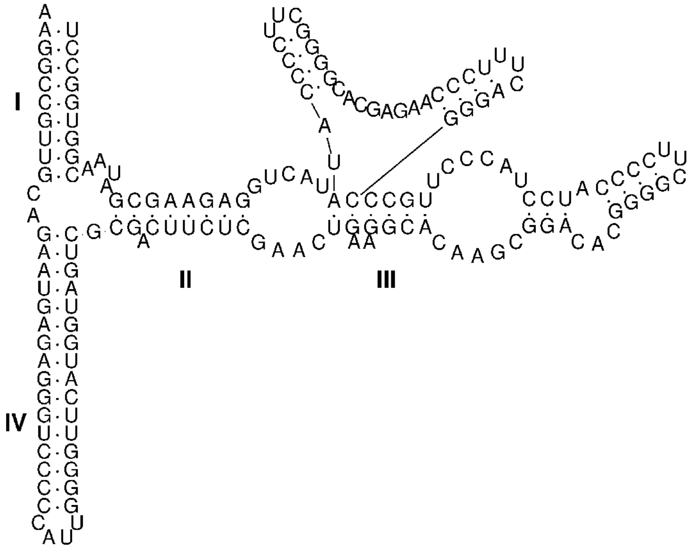


**Desulfofarcimen acetoxidans 5S rRNA**
**NC_013216.1[272527-272641]fwd**

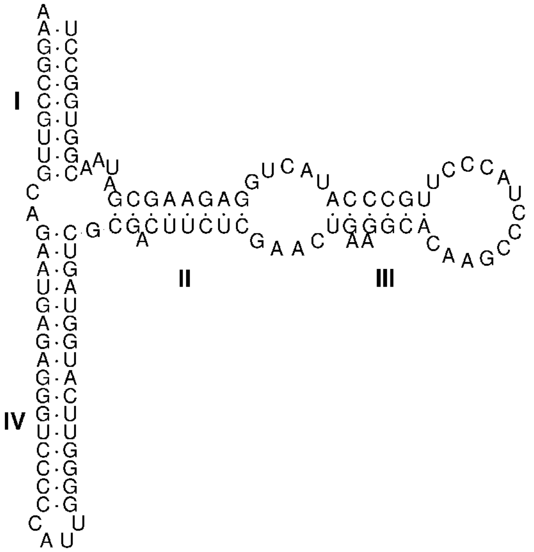


**Supplementary Figure 2**. Variant 5S rRNAs with or without expansion segments identified in *Desulfofarcimen acetoxidans* DSM 771 genome.
