## Supplemental Fig. S3 for "Expansion Segments in Bacterial and Archaeal 5S Ribosomal RNAs"

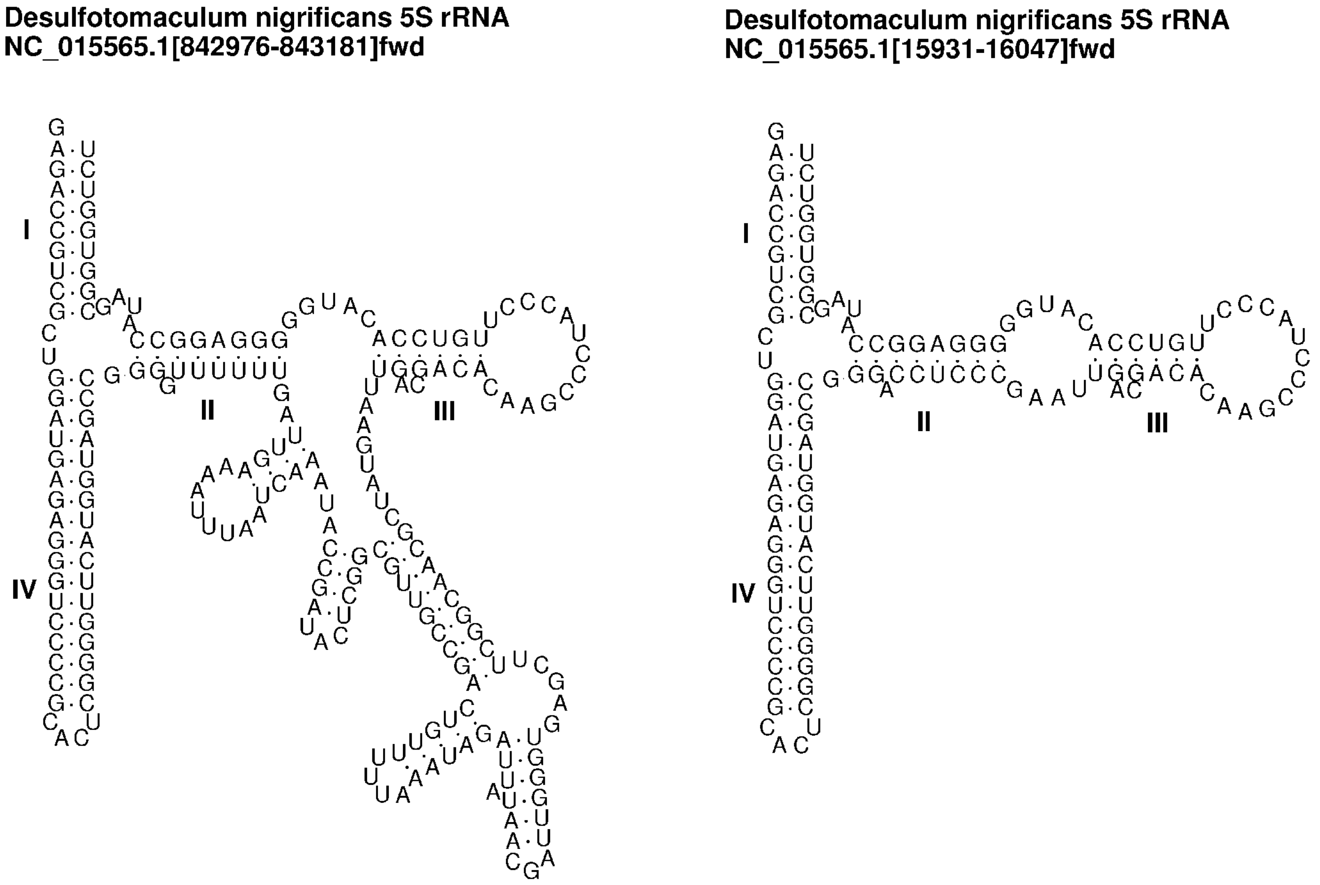


**Supplementary Figure 3**. Variant 5S rRNAs with or without expansion segments identified in *Desulfotomaculum nigrificans* CO-1-SRB genome.
