## Supplemental Fig. S4 for "Expansion Segments in Bacterial and Archaeal 5S Ribosomal RNAs"

**Desulfotomaculum reducens 5S rRNA**
**NC_009253.1[361111-361315]fwd**

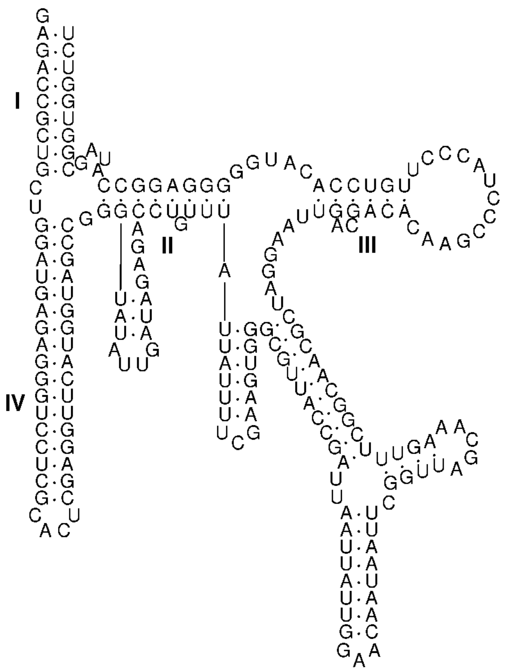


**Desulfotomaculum reducens 5S rRNA**
**NC_009253.1[2219507-2219711]rev**

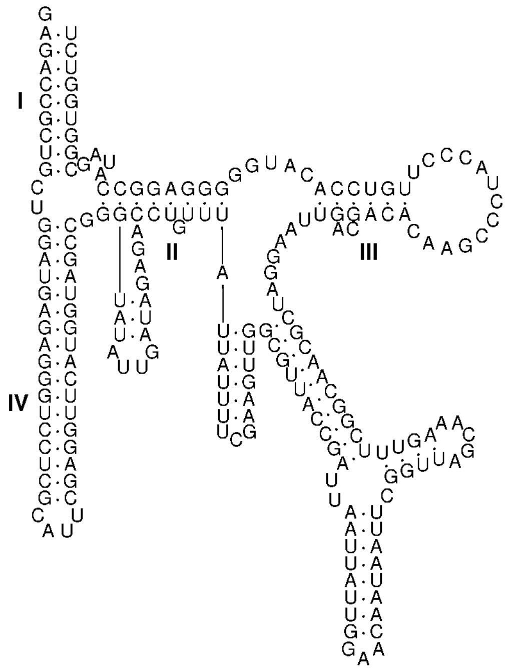


**Desulfotomaculum reducens 5S rRNA**
**NC_009253.1[15845-15961]fwd**

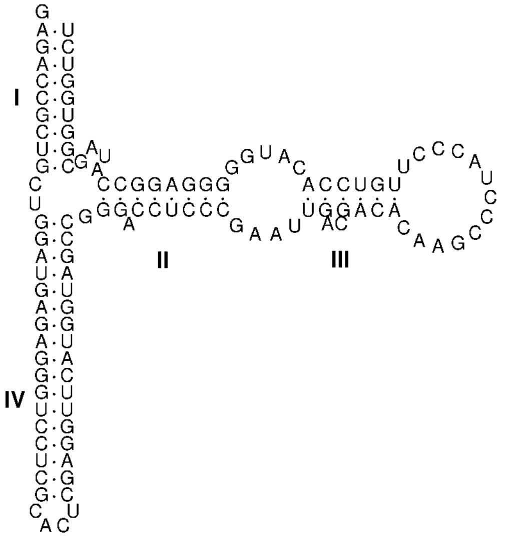


**Supplementary Figure 4**. Variant 5S rRNAs with or without expansion segments identified in *Desulfotomaculum reducens* MI-1 genome.
