## Supplemental Fig. S5 for "Expansion Segments in Bacterial and Archaeal 5S Ribosomal RNAs"

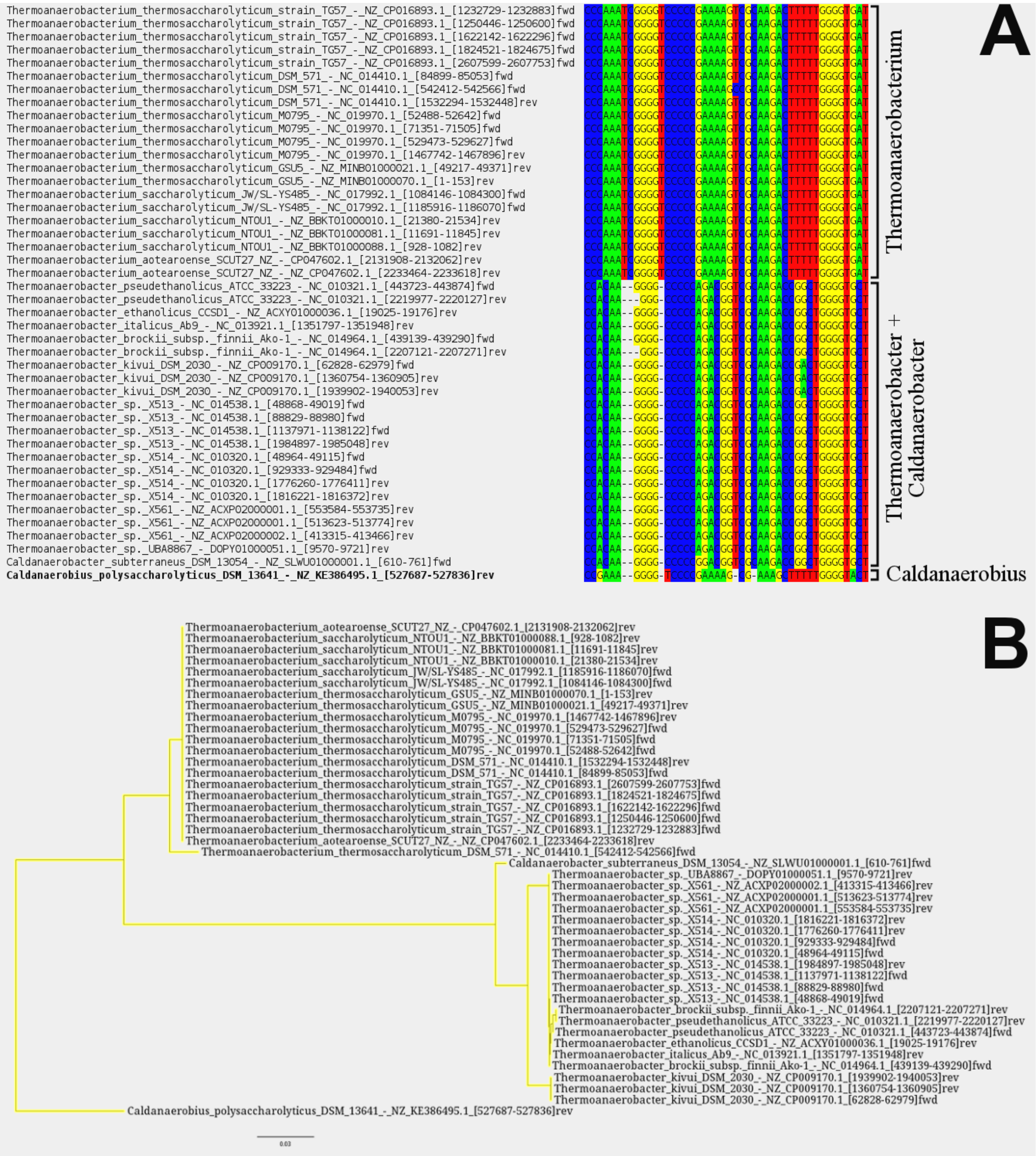


**Supplementary Figure 5**. Multiple sequence alignment of the expansion segments identified in 5S rRNAs from four thermoanaerobacterial genera, *Thermoanaerobacterium*, *Thermoanaerobacter*, *Caldanaerobacter*, and *Caldanaerobius* (**A**), and corresponding Neighbor Joining tree (**B**).
