## Supplemental Fig. S6 for "Expansion Segments in Bacterial and Archaeal 5S Ribosomal RNAs"

### Covariation analysis of predicted secondary structure of the expansion segments found in halobacterial 5S rRNAs

A. R-scape consensus 2D structure

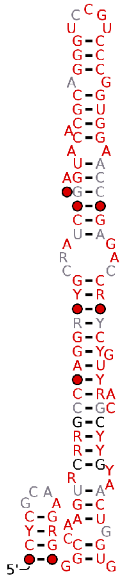

B. RNAalifold consensus 2D structure

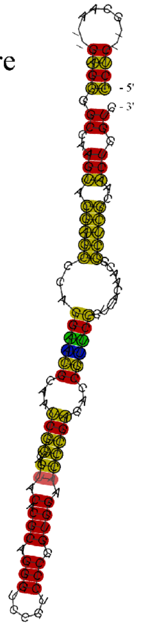

C. RNAalifold multiple RNA sequence alignment

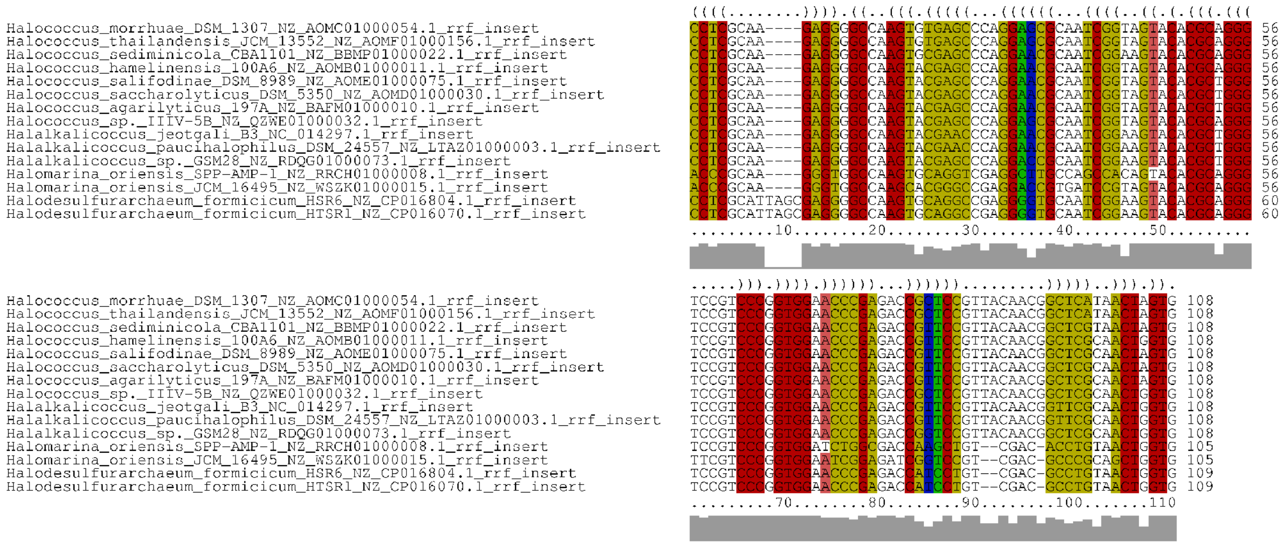


**Supplementary Figure 6**. Covariation analysis of predicted secondary structure of the expansion segments identified in 5S rRNAs from four halobacterial genera, *Halococcus*, *Halalkalicoccus*, *Halomarina*, and *Halodesulfurarchaeum*.
