## Supplemental Fig. S7 for "Expansion Segments in Bacterial and Archaeal 5S Ribosomal RNAs"

Covariation analysis of predicted secondary structure of the expansion segments found in thermoanaerobacterial 5S rRNAs

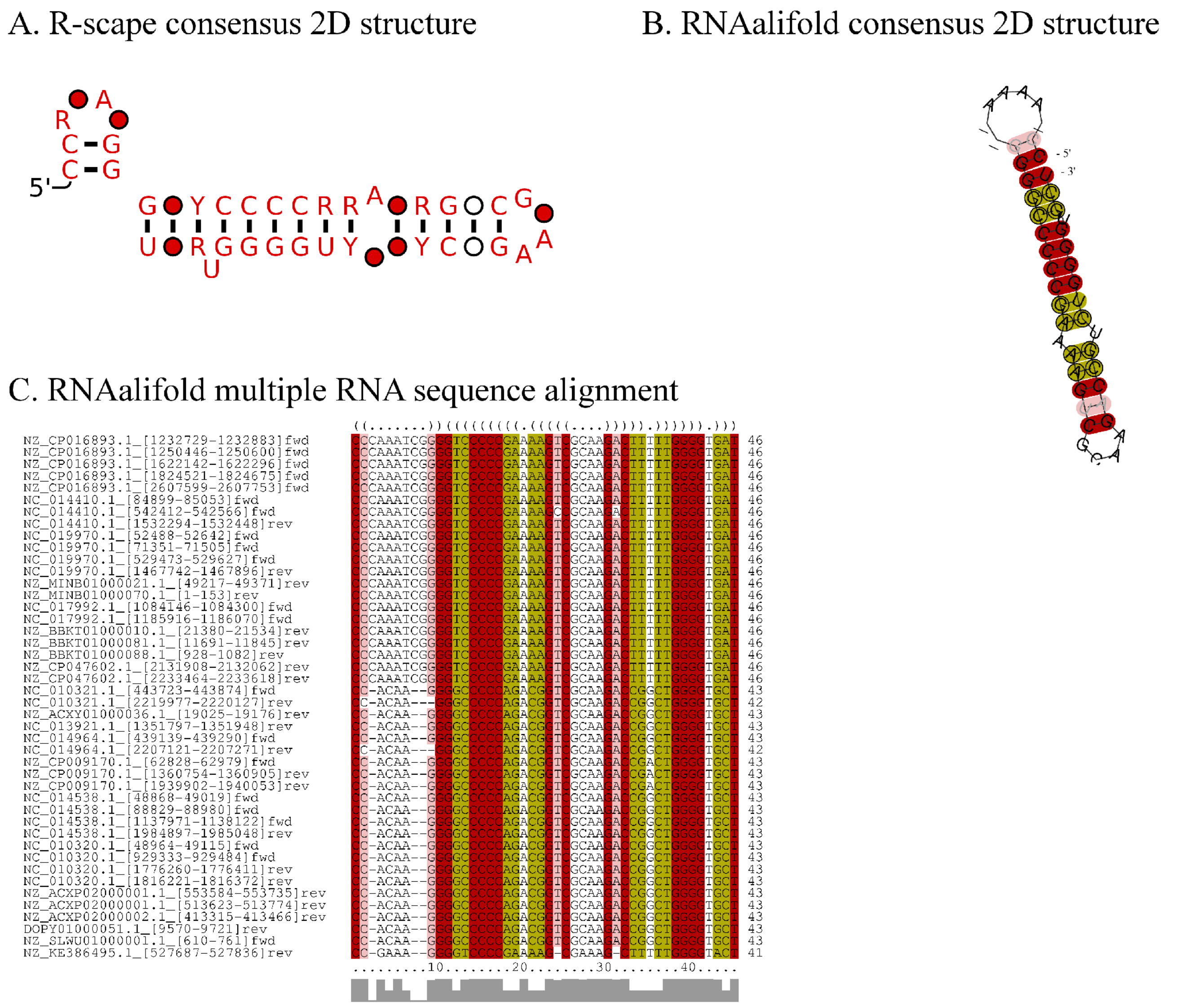


**Supplementary Figure 7**. Covariation analysis of predicted secondary structure of the expansion segments identified in 5S rRNAs from four thermoanaerobacterial genera, *Thermoanaerobacterium*, *Thermoanaerobacter*, *Caldanaerobacter*, and *Caldanaerobius*.
