## Supplemental File S1 for "Expansion Segments in Bacterial and Archaeal 5S Ribosomal RNAs": description.txt

Supplementary File S1. Sequences of all identified genes of 5S rRNAs with expansion segments.

The multi-entry FASTA file contains 89 non-aligned sequences of the expanded 5S rRNA genes from
the archaeal order Halobacterales and bacterial orders Clostridiales, Halanaerobiales,
Thermoanaerobacterales, and Alteromonadales.
