## Supplemental File S2 for "Expansion Segments in Bacterial and Archaeal 5S Ribosomal RNAs": description.txt

Supplementary File S2. Collection of multiple sequence alignments of valid 5S rRNA genes
and putative 5S rRNA pseudogenes from several representatives of the bacterial
families Peptococcaceae and Pseudoalteromonadaceae.

Each file contain multiple alignment of sequences originating from the same organism.
The alignments are in FASTA format. Each putative 5S rRNA pseudogene situated downstream of
of a particular valid 5S rRNA gene is designated as a "satellite" of this gene. Stand-alone 5S
rRNA pseudogenes are designated as "remnants".
