## Supplemental File S3 for "Expansion Segments in Bacterial and Archaeal 5S Ribosomal RNAs": description.txt

Supplementary File 3. Collection of multiple sequence alignments of 16S, 23S, and 5S rRNAs from
several representatives of the bacterial genera Desulfallas, Desulfitobacterium, Desulfofarcimen,
Thermoanaerobacter, and Thermoanaerobacterium. Each of the presented organisms carries at least one
expanded 5S rRNA gene and at least one expanded gene of 16S or 23S rRNAs.

Each file contain multiple alignment of sequences of the same type originating from the same organism.
rRNA types are indicated by suffixes 16s, 23s, and 5s, respectively. The alignments are in FASTA format.
